# Neural correlates of multisensory enhancement in audiovisual narrative speech perception: a fMRI investigation

**DOI:** 10.1101/2022.02.14.480408

**Authors:** Lars A. Ross, Sophie Molholm, John S. Butler, Victor A. Del Bene, John J. Foxe

## Abstract

This fMRI study investigated the effect of seeing articulatory movements of a speaker while listening to a naturalistic narrative stimulus. It had the goal to identify regions of the language network showing multisensory enhancement under synchronous audiovisual conditions. We expected this enhancement to emerge in regions known to underlie the integration of auditory and visual information such as the posterior superior temporal gyrus as well as parts of the broader language network, including the semantic system. To this end we presented 53 participants with a continuous narration of a story in auditory alone, visual alone, and both synchronous and asynchronous audiovisual speech conditions while recording brain activity using BOLD fMRI. We found multisensory enhancement in an extensive network of regions underlying multisensory integration and parts of the semantic network as well as extralinguistic regions not usually associated with multisensory integration, namely the primary visual cortex and the bilateral amygdala. Analysis also revealed involvement of thalamic brain regions along the visual and auditory pathways more commonly associated with early sensory processing. We conclude that under natural listening conditions, multisensory enhancement not only involves sites of multisensory integration but many regions of the wider semantic network and includes regions associated with extralinguistic sensory, perceptual and cognitive processing.

## INTRODUCTION

Sampling of information through multiple sensory systems is of fundamental utility to an organism since it enhances the likelihood of both detection and identification of survival-relevant objects or events in the environment. Initially channeled separately, different sensory inputs pertaining to the same objects or events are integrated across multiple stages of sensory and perceptual processing, leading to enhancements of behavior such as improved accuracy and faster reaction times for perceptual judgments (Bolognini et al., 2007; Brandwein et al., 2014; Brandwein et al., 2011; Diederich & Colonius, 2004; Foxe & Molholm, 2009; Frens et al., 1995; Molholm et al., 2004; Molholm et al., 2002; Nozawa et al., 1994; Rowland et al., 2007; Sperdin et al., 2009; Stein et al., 1989). Multisensory integration (MSI) organizes and reduces the complexity of our sensory environment by binding multiple sensory inputs into single, unified percepts. It has been hypothesized that a failure of this function can lead to a sensory environment that is perceived as overwhelming with potential consequence of perceptual and behavioral deficits and maladaptive responses toward the environment (Ayres, 1979; Brandwein et al., 2015; Foxe & Molholm, 2009; Molholm et al., 2020).

One area of particular interest to multisensory (MS) researchers is speech recognition, where it has long been known that visual articulatory cues can strongly influence auditory speech perception (McGurk & MacDonald, 1976; Saint-Amour et al., 2007; Tjan et al., 2014). This is especially true when the auditory speech signal is ambiguous, as is often the case when the background environment is noisy or there are multiple simultaneous speakers (Benoit et al., 1994; Foxe et al., 2020; Foxe et al., 2015; Ma et al., 2009; MacLeod & Summerfield, 1987; Molholm et al., 2020; Richie & Kewley-Port, 2008; Ross et al., 2011; Ross, Saint-Amour, Leavitt, Javitt, et al., 2007; Senkowski et al., 2008; Sumby, 1954). Indeed, despite the fact that most of us are generally poor lip readers when only the visual signal is available, the enhancing effects of visual speech can be dramatic, such that visual inputs can render completely indecipherable vocalizations clearly audible.

There is a great deal of disagreement about the mechanisms of speech processing (Navarra, 2012) such as the format of the representations underlying the perception of speech (Massaro, 1987; Stevens, 2002; Liberman and Mattingly, 1985). This, in turn poses challenges to neuroscientific models of audiovisual speech processing, particularly where and when information from different sensory channels is integrated in the brain.

A common approach to investigating the neural mechanisms of AV speech processing is to use neuroimaging to compare hemodynamic responses to MS speech with responses to the constituent unisensory components (i.e., AV speech vs. auditory-alone or visual-alone speech). This allows for isolation of neural regions that show stronger responses to AV speech. The region most consistently localized is the superior temporal sulcus/gyrus (pSTS/G), an area well-known for its involvement in AV integration. This is the case whether the stimuli are as simple as nonsense monosyllables (Callan et al., 2003; Okada et al., 2013; Reale et al., 2007) and single words (Calvert et al., 1999; Wright et al., 2003), or as complex as a spoken story (Calvert et al., 2000). The MSI role of the pSTS/G extends to other aspects of AV speech stimuli such as congruency (Murase et al., 2008; Nath & Beauchamp, 2011), temporal synchrony (Macaluso et al., 2004; Noesselt et al., 2012), and ambiguity (Saint-Amour et al., 2007; Sekiyama et al., 2003; Stevenson & James, 2009). Other regions such as the primary auditory cortex (Calvert et al., 1999) and the motor cortex (Schomers & Pulvermuller, 2016) have also been implicated in AV speech processing suggesting more than one mechanism underling observed MSI effects (Navarra, 2012). However, these areas have not been reliably implicated across studies and have received relatively little attention. The lack of consistency across studies regarding the regions that comprise the wider MSI speech network likely reflects, at least in part, the use of different paradigms, different stimulus materials and the use of different criteria to assess MS integration in the BOLD signal. However, we suspect that a major reason for between-study variability is simply that the studies often have relatively low levels of statistical power due to modest sample sizes. Most of the AV speech processing studies discussed so far are based on sample sizes below 15 (except N = 28 in the study of (Murase et al., 2008) and N = 20 in the study of (Okada et al., 2013)). A notable exception is a large-scale lesion study (*N* = 100) on AV integration in speech (Hickok et al., 2018).

Small sample sizes in fMRI studies represent a potentially serious limitation because different MS regions are engaged in different processes, and activity in these regions is likely to be recruited with varying strengths. For example, in cases where some nodes of the MS network are engaged in a relatively transient manner and some regions receive modulatory rather than direct afferent inputs, effects in these regions would be expected to be modest (Allman & Meredith, 2007) and would likely go undetected in studies with modest sample sizes. The present study sought to significantly increase sensitivity to smaller effect sizes by testing a considerably larger sample than is typical of this type of study (N = 53), with a main goal of reliably characterizing the network of brain regions involved in AV speech processing.

The strong consensus regarding involvement of pSTS/G is based on highly stringent criteria. That is, this region survives many different types of experimental manipulations and statistical analysis methods and criteria across the many studies to examine assorted aspects of AV speech processing (Beauchamp, 2005; Calvert, 2001). However, speech is a complex stimulus and its processing engages a widely distributed network of regions serving a broad range of functions from sensory to semantic processing (Hickok & Poeppel, 2007; Price, 2010; Rauschecker, 2012). As such, the visual benefit manifested in enhanced speech perception is highly unlikely to be related solely to the involvement of a single region (i.e., pSTS/G), but rather, must surely arise from coordinated activity across the network of speech processing regions. It has been shown that MS interactions occur at multiple stages of information processing (Foxe & Schroeder, 2005) and thus, AV speech processing should be reflected in activity of multiple regions. Indeed, a number of studies have reported that AV speech amplifies activity in regions as early in the information processing hierarchy as the primary auditory cortex (Callan et al., 2003; Calvert et al., 1999; Calvert et al., 1997; Calvert et al., 2000; Okada et al., 2013), visual motion regions (Puce et al., 1998; Puce et al., 2003; Wright et al., 2003; Yarkoni et al., 2011), and prefrontal regions such as Broca’s area and premotor cortex (Iacoboni, 2008; Meister et al., 2007; Ojanen et al., 2005; Skipper et al., 2005; Wilson et al., 2004).

Further, it is reasonable to assume that under natural listening conditions, integration in MS regions has downstream consequences in the larger speech and language network. For example, if AV integration results in improved perception of the auditory speech signal, then the consequences of integration should be observable in the BOLD response in regions underlying the perception and semantic processing of the respective speech stimulus at word, sentence and narrative levels. Further, it may be possible, depending on the content, to observe enhancing effects on other cognitive functions such as memory retrieval and emotional processing. Since most studies investigating MS integration are interested in the processes and regions underlying MS modulation and convergence, these effects have been considered confounds with the intention to eliminate them through experimental control. However, this deprives us of the observation of broader effects that AV integration might have on the speech processing and cognitive network. Moreover, most studies investigating AV integration used truncated speech material such as syllables and words often in the context of a McGurk-type paradigm where participants are asked to reconcile conflicting auditory and visual cues. It has been questioned whether these tasks engage the same mechanisms active in more natural AV speech processing (Alsius et al., 2018; Hickok et al., 2018; Peelle, 2019; Van Engen & Peelle, 2014). We therefore expect these broader MS enhancement effects to arise with natural and more complex stimulus material such as narratives. For the purpose of this investigation, we use the term MS enhancement in a broader sense to refer to processes of MS integration and their possible consequences on linguistic and cognitive processing because our experimental approach does not strictly distinguish them from one another.

Therefore, the present study had several central goals. The first was to comprehensively map the network of brain regions involved in AV speech perception in a large sample of healthy adults. Importantly, this study focused not only on identifying regions of AV integration but also aimed to assess the downstream effects of AV integration on the larger language network engaged when listening to natural narrative speech.

Using BOLD fMRI, we presented continuous natural speech in varying conditions: auditory alone (A), visual alone (V), synchronous audiovisual (AV) and an asynchronous audiovisual condition (AVa). We characterized regions engaged in the presentation of the unisensory conditions (A and V) in order to identify brain regions engaged in natural narrative speech processing and speechreading respectively. We mapped AV enhancement effects by examining areas that responded more strongly to the AV speech compared to unisensory speech stimuli. We employed the maximum criterion (Beauchamp, 2005; James, 2012) by performing a conjunction analysis identifying regions in which the AV-response was significantly larger than the A and the V response [(AV > A) ⍰ (AV > V)] while constraining the analysis to regions in which AV was significantly larger than baseline (AV > 0). We also explored which regions of the identified network showed a superadditive response to the AV stimulus [AV > (A + V)].

To further consider the regions key to the processing of MS speech, we added an experimental condition where the AV inputs were out of synchrony and compared responses to synchronous and asynchronous AV speech. The purpose was to investigate regions sensitive to the temporal alignment of the AV speech signals, under the assumption that one way that MSI occurs is via binding of different sensory signals that are correlated in time (Stein et al., 1988).

In a final exploratory analysis, we assessed the relationship between activation to the respective conditions in the fMRI experiment and behavioral measures on an AV speech perception task obtained from the same subjects in an experiment completed outside the scanner. The goal was to test for relationships between AV-integration performance and the BOLD response to identify regions associated with audio-visual enhanced speech perception.

## MATERIALS AND METHODS

### Participants

From an original sample of 60 participants, 7 were excluded based on technical difficulties during the scan or post processing, due to excess motion or lack of task compliance. The data of 53 native English-speaking adults with no history of neurological and psychiatric problems and no substance abuse (25 female, age range = 20 years of age to 35 years of age, *M* = 25 years, *SD* = 3.8 years) were included in the following fMRI analyses. All had normal hearing and normal or corrected-to-normal vision. Out of the 53, 47 were right-handed, 3 left-handed and 2 were ambidextrous (Oldfield, 1971). Handedness of one participant was not recorded. The study was approved by the Institutional Review Board of the Albert Einstein College of Medicine and all procedures were conducted in accordance with the tenets of the Declaration of Helsinki. All participants gave written informed consent and were paid for their participation.

### MRI acquisition

Imaging data were acquired using a 3.0 Tesla Philips Achieva TX scanner with a 32-channel head coil. A T1-weighted whole-head anatomical volume was obtained using a 3D magnetization-prepared rapid gradient-echo (MP-RAGE) sequence (echo time [TE] = 3.7 ms, repetition time [TR] = 8.2 ms, flip angle [FA] = 8 degrees, voxel size = 1 x 1 x 1 mm^3^, matrix = 256 x 256, FOV = 256 x 256 mm^2^, number of slices = 220). T2*-weighted functional scans were acquired using gradient echo-planar imaging (EPI). This acquisition covered the whole brain excluding inferior aspects of the cerebellum below the horizontal fissure (axial acquisition in ascending order, TE = 20 ms, TR = 2000 ms, FA = 90 degrees, voxel size = 1.67 x 1.67 x 2.30 mm^3^, matrix = 144 x 144, FOV = 240 x 240 mm^2^, number of slices per volume = 50, total number of volumes = 158 + 172 + 146).

### fMRI task

Participants were presented with video recordings of a speaker reading from a children’s story about economic and environmental issues called “The Lorax” written by Dr. Seuss. The video of the story (lasting 14 min 38 s) was segmented into sections of varying length ranging from 8 to 22 s. The frame rate of the video recordings was 29 frames per second. Each section was randomly assigned for each participant to one of four conditions: auditory (A), visual (V), synchronous audiovisual (AV), and asynchronous audiovisual (AVa). As such, block length is a random variable that is not associated with a given condition. The A and V conditions presented the auditory and visual stimuli alone, respectively. During the A condition, a still image of the speaker was presented and participants were told to look at the picture while listening to the story. The AV and AVa conditions presented both the auditory and visual stimuli, but in the AV condition, the two inputs were presented in synchrony whereas in the AVa conditions, the visual input was delayed by 400 ms relative to the timing of the auditory input such that the audio and video were clearly misaligned. The story was presented in 3 runs of 4 min 50 s, 5 min 20 s, and 4 min 28 s, respectively, with each run presenting a sub-story in a continuous manner. Participants were instructed to follow the whole story carefully regardless of the changing presentation mode. The story in each run was followed by a resting period during which a screen containing a sign saying “please relax” was presented briefly and disappeared, leaving only a blank screen. Participants were asked to rest during this period with their eyes open. The resting period lasted 18, 16 and 16 s for the respective 3 runs. There were no rest periods between blocks and for a given contrast the baseline represents the average time course. Retention of the story content was assessed with a 10-item story comprehension questionnaire after the scan.

Throughout the whole MRI session, participants wore foam ear plugs to attenuate the scanner noise and MR-compatible headphones (the Serene Sound system; Resonance Technology, Inc.) through which the auditory stimuli were presented. The SPL of the headphones was kept constant in the range of 90 to 95 dB across the participants who reported this volume to be audible and comfortable. Participants wore MR-compatible glasses (the VisuaStim Digital system; Resonance Technology, Inc.) through which the visual stimuli were delivered at a refresh rate of 60 Hz. An eye tracker (the MReyetracking system; Resonance Technology, Inc.) was mounted inside the glasses and used to monitor that participants’ eyes were open and watching the video, throughout the task.

### fMRI analysis

All imaging data were analyzed in BrainVoyager (version 22.2, Brain Innovation, Maastricht, the Netherlands). The functional data were pre-processed using interscan slice time correction (cubic spline interpolation) and 3D rigid-body motion correction (trilinear sinc interpolation). The data of all three runs were aligned to the first volume of the first run. No subject data were removed for excess motion based on a cutoff of 2mm/degrees in any direction). Individual anatomical images were transformed into Talairach space (sinc interpolation) and functional imaging data were aligned to the individual’s anatomy using boundary-based registration (Greve & Fischl, 2009) and inspected for quality of registration. The time courses for each participant were subsequently temporal high pass filtered with a GLM Fourier basis set and spatially smoothed using a 6mm FWMH Gaussian Kernel before transformation into Talairach space.

Voxel-wise statistical analyses were performed on the (%) normalized functional data using a two-level random-effects GLM approach with A, V, AV and AVa as predictors which were convolved with a standard two-gamma hemodynamic response function. We used a Talairach mask to exclude voxels outside the brain.

The following contrasts of interest and conjunction analyses were performed and reported: (1) A vs baseline: This analysis was performed to identify brain regions active during the processing of the story (narrative) without visual articulatory information. (2) V vs baseline: Here, we investigated regions involved in the processing of visual articulatory information. (3) [(AV-A) ⍰ (AV-V)]: This was the analysis critical for the identification of MS enhancement according to the Max Criterion (Beauchamp, 2005; James, 2012) and tests via mathematical conjunction of the (AV-A) and (AV-V) contrasts (Nichols et al., 2005) whether activation to the AV condition significantly supersedes the A and the V condition against their baseline. In regions meeting this criterion activation to the AV condition is significantly larger than activation to the A condition *and* activation to the V condition (4) (AV > A+V): We also tested AV enhancement according to the additive (superadditive) criterion (Calvert & Thesen, 2004) where the BOLD response to the AV condition was larger than the sum of the A and V responses. For this analysis we summed the normalized predictor values for the A and V conditions from each voxel and subtracted them from the predictor values of the AV condition. The resulting values were tested against zero using a t-test. (5) AV vs. AVa: In this contrast we compared the synchronous V and the asynchronous AV condition.

The following analyses were secondary in regard to the goals of this study and are reported in the appendix: (6) A vs V: The difference between auditory and visual conditions. This contrast is particularly sensitive to activations in the auditory and visual cortices and was applied on a single subject basis after a fixed effects GLM with the predictors of interest to assure the compliance to the experimental instructions and the absence of failed data acquisition due to technical problems. On a group level, this analysis was performed to delineate regions where both conditions differed from one another and allow a comparison to the statistical map of regions where they were active in conjunction, as follows. (7) A ⍰ V: The conjunction of the contrasts (Nichols et al., 2005) of A and V conditions against baseline tests for voxels in which both A and V conditions differ significantly from baseline. We were interested in this analysis primarily to determine whether regions of MS enhancement are also responsive to the A and V conditions.

For all whole brain analyses we used the false discovery rate (FDR) procedure (Genovese et al., 2002) to control for multiple comparisons at q < 0.05.

### Out of scanner MS speech recognition behavioral task

Stimulus materials consisted of digital recordings of 300 simple monosyllabic words spoken by a female speaker. This set of words was a subset of the stimulus material created for a previous experiment in our laboratory (Ross, Saint-Amour, Leavitt, Javitt, et al., 2007) and used in a previous study (Ross et al., 2011). These words were taken from the “MRC Psycholinguistic Database” (Coltheart, 1981) and were selected from a well-characterized normed set based on their written-word frequency (Kucera & Francis, 1967). The subset of words for the present experiment is a selection of simple, high-frequency words likely to be in the lexicon of participants in the age-range of our sample. The recorded movies were digitally re-mastered so that the length of the movie (1.3 sec) and the onset of the acoustic signal were similar across all words. Average voice onset occurred at 520ms after movie onset (SD= 30ms). The words were presented at approximately 50dBA FSPL, at seven levels of intelligibility including a condition with no noise (NN) and six conditions with added pink noise at 50, 53, 56, 59, 62 and 65dBA FSPL sound pressure. Noise onset was synchronized with movie onset. The signal-to-noise ratios (SNRs) were therefore NN, 0, −3, −6, −9, −12, −15dBA FSPL. These SNRs were chosen to cover a performance range in the auditory-alone condition from 0% recognized words at the lowest SNR to almost perfect recognition performance with no noise. The movies were presented on a monitor (NEC Multisync FE 2111SB) at 80cm distance from the eyes of the participants. The face of the speaker extended approximately 6.44° of visual angle horizontally and 8.58° vertically (hairline to chin). The words and pink noise were presented over headphones (Sennheiser, model HD 555).

The main experiment consisted of three randomly intermixed conditions: In the auditory-alone condition (A-alone) the auditory words were presented in conjunction with a still image of the speakers face; in the AV condition the auditory words were presented in conjunction with the corresponding video of the speaker articulating the words. Finally, in the visual alone condition (V-alone) only the video of the speaker’s articulations was presented. The word stimuli were presented in a fixed order and the condition (the noise level and whether it was presented as A-alone, V-alone or AV) was assigned to each word randomly. Stimuli were presented in 15 blocks of 20 words with a total of 300 stimulus presentations. There were 140 stimuli for the A and AV conditions respectively (20 stimuli per condition and intelligibility level) and 20 stimuli for the V condition that was presented without noise.

Task: Participants were instructed to watch the screen and report which word they heard (or saw in the V-alone condition). If a word was not clearly understood, participants were encouraged to make their best guess. An experimenter, seated approximately 1 m distance from the participant at a 90° angle to the participant-screen axis, monitored participant’s adherence to maintaining fixation on the screen. Only responses that exactly matched the presented word were considered correct. Any other response was recorded as incorrect.

#### Analyses of Task Performance

We submitted percent correct responses for each condition to a repeated measures analysis of variance (RM-ANOVA) with factors of stimulus condition (A vs. AV), SNR level (7 levels) and biological sex as well as age as covariates. Performance in the Valone condition was analyzed separately because it was only presented without noise. Violations of the sphericity assumption of the RM-ANOVA were corrected by adjusting the degrees of freedom with the Greenhouse-Geisser correction method. MS enhancement (or AV-gain) was operationalized here as the difference in performance between the AV and the A-alone condition (AV – A-alone). This analysis was performed at the four lowest SNRs because the variance at higher SNRs becomes increasingly constrained by ceiling performance (Ross et al., 2011). We performed two-tailed Pearson correlation tests between the A and the AV conditions at the four lowest SNRs (average) to determine if A-performance under noisy conditions was negatively associated with AV-performance at the same SNRs. We also tested for an association between the A and V conditions.

Finally, we also tested the hypothesis that individuals with more difficulty perceiving auditory speech when it is masked with noise are better speech-readers, who therefore benefit more from AV input. The presence of this trade-off gained recent support from a study showing that early electrophysiological indices of auditory processing predict auditory, visual and AV speech processing (Dias et al., 2021). If such effects were apparent in our data, our goal was to investigate possible relationships of auditory processing ability with measures of brain activity in our fMRI study

#### Correlation with BOLD measures

For the analysis of MS enhancement we averaged MS gain at the four lowest SNRs where most audiovisual enhancement was observed and computed voxel-wise correlations with the beta weights of the AV-A contrast of our BOLD data. The resulting correlation (Pearson’s r) maps were thresholded at *p* = 0.001 as suggested by recent findings (Eklund et al., 2016) in order to control for family wise error rate (FWE). If this initial threshold produced a map lacking a sufficient size and distribution of significant clusters, this threshold was iteratively increased to a maximum of *p* = 0.01. This compromise in regard to the risk for false positives was motivated by our expectation that the effect size of the correlation of our GLM predictors with measures of behavioral performance would not be of the same magnitude as the effect size of moderate BOLD effects for which a *p* < 0.001 threshold was shown to be appropriate. We therefore did not expect this threshold would result in a statistical map with a cluster distribution suitable for a subsequent Monte Carlo simulation. We used the thresholded map as input for the Cluster-Level Statistical Threshold Estimator plugin in Brainvoyager using 5000 iterations. This tool simulates the distribution of normally distributed noise based on the smoothness of the map used as input in each iteration step and records the frequency and size of the resulting clusters. We performed exploratory analyses of the relationship between BOLD effects and behavioral performance in the A, V and AV conditions.

##### Association between questionnaire performance and BOLD measures

Finally, we explored relationships between questionnaire performance and BOLD measures. There are multiple possible variables affecting questionnaire performance including the quality of recall at the time of test administration, improper comprehension of the story due to lack of attention to the stimulus, varying stimulus delivery or cognitive ability. While we are not able to resolve the precise cause of performance variability we can assess if mechanisms giving rise to the variability are observable as brain activity during the scan. The location (eg. sensory, perceptual or semantic regions) of these BOLD effects may favor one or more possible causes over others. If sufficient variability in questionnaire performance is observed we intended to explore relationships of this measure with brain activity to the A and AV conditions.

## RESULTS

### Auditory alone (A)

The stimulation in the auditory alone condition (A vs. baseline) resulted in strong activation in bilateral Heschl’s gyrus with peak activations within the primary auditory cortex (see Fig, 1 and Table 1). From these locations, clusters in both hemispheres extended anteriorly along the superior temporal gyrus and its upper bank into the anterior temporal lobes (ATLs) including a cluster in the ventral ATL in the left hemisphere, laterally along the transverse temporal gyrus and posteriorly toward the posterior parietal junction. Activations extended from the primary auditory cortices into the ventral motor cortex along the roofs of the lateral sulci covering the parietal operculae in both hemispheres. The auditory condition also engaged left hemispheric regions in the frontal lobe including the inferior frontal gyrus (IFG), the dorsolateral prefrontal cortex (dlpfc). The supplementary motor cortex was engaged in both hemispheres.

**Figure 1.**
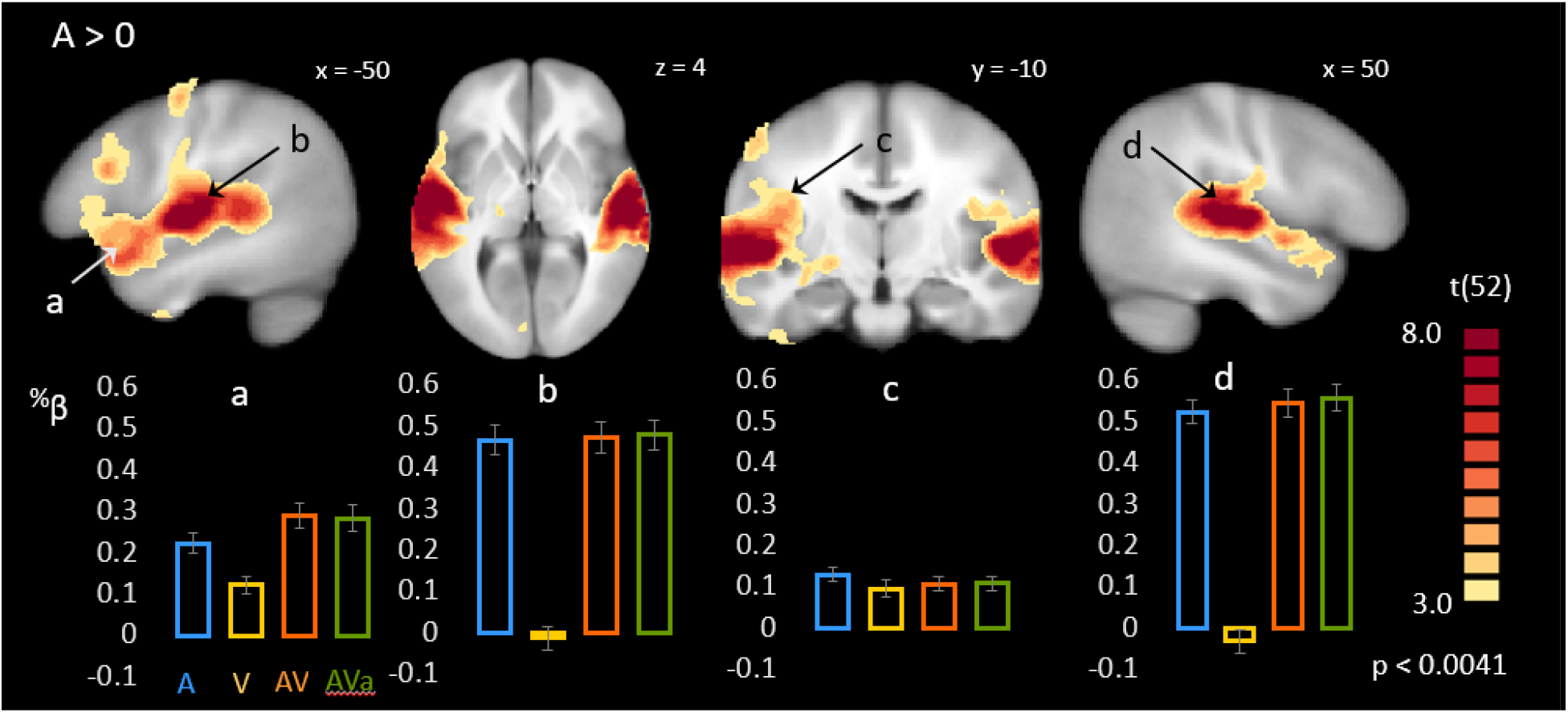
Statistical comparison of the A condition to baseline. Maps show voxels with significant t-scores of the comparison of the A-condition to baseline FDR-corrected (q = 0.05) for multiple comparisons. Bar graphs represent selected % transformed predictor values for A, V, AV and AVa conditions averaged over 4 functional voxels centered around peak voxel locations (see Table 1). a) Left anterior superior temporal gyrus; b) left Heschl’s gyrus; c) left insula; d) right Heschl’s gyrus.

**Table 1.**
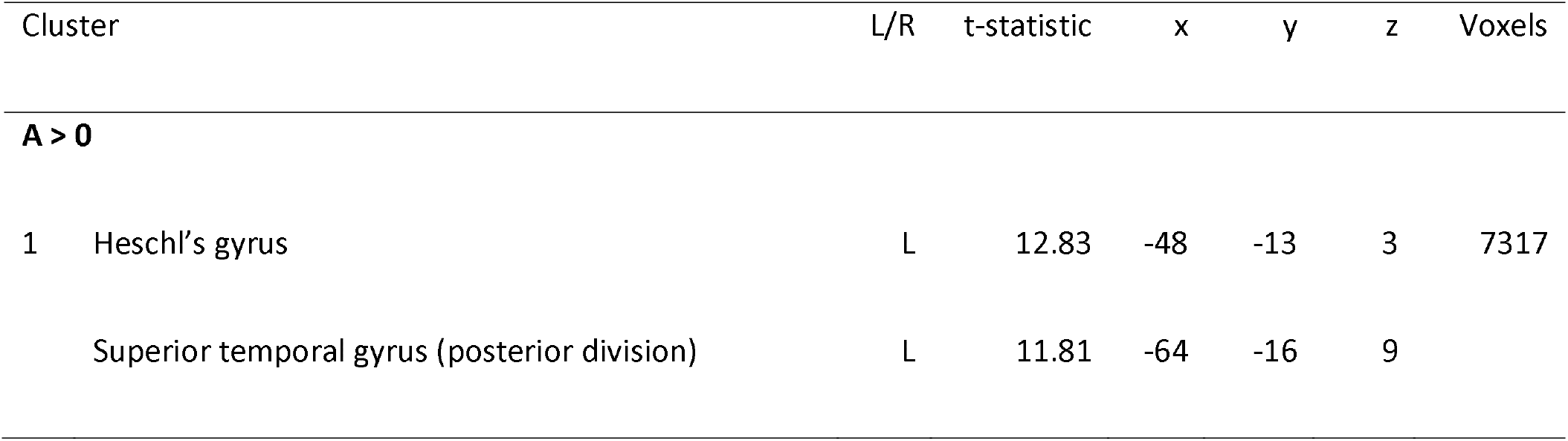
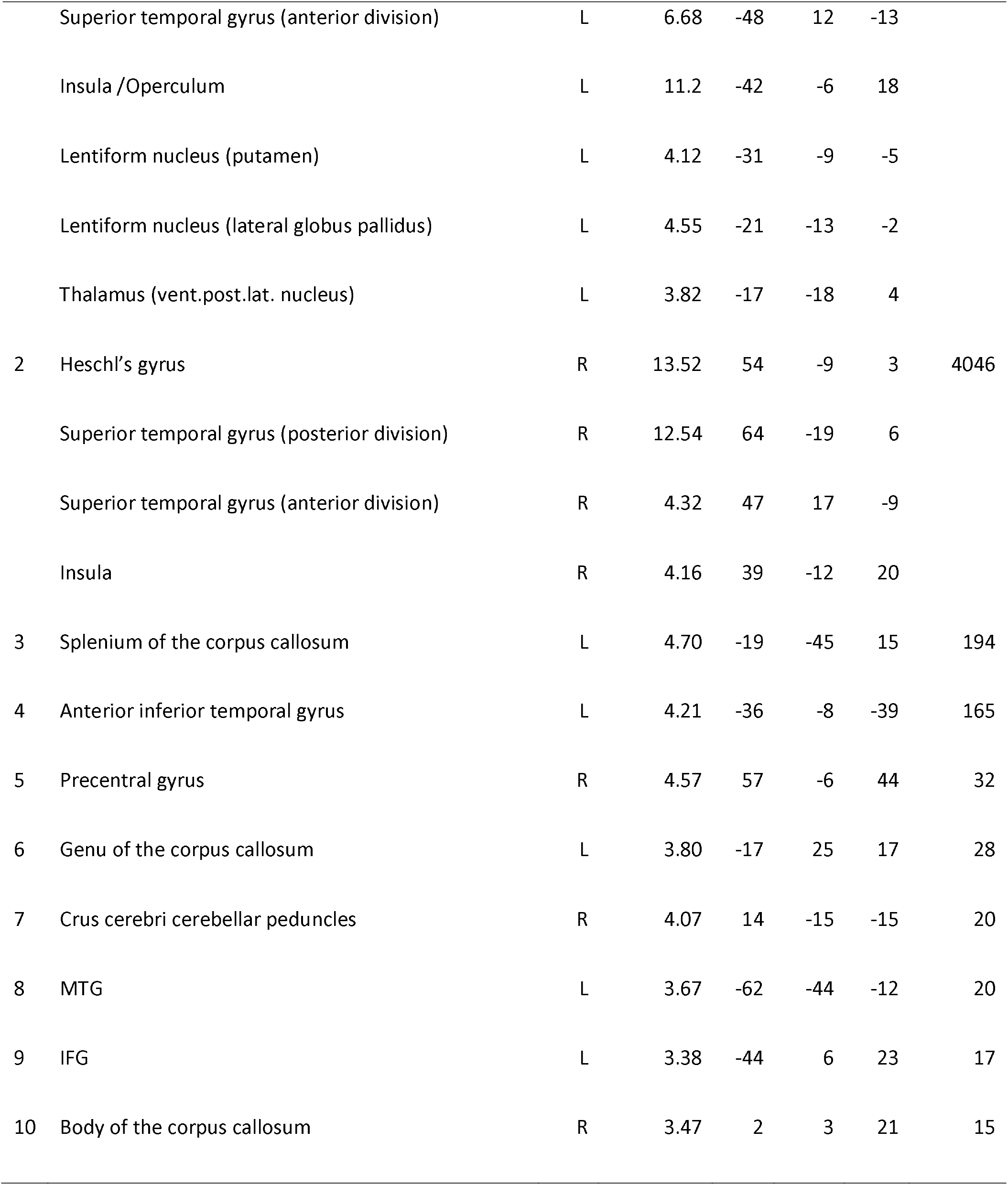
Clusters of significant activity resulting from the contrast of the A condition vs. baseline. Significant clusters are numbered and reported with their t-statistic and location in Talairach space in the order of cluster size. In cases where clusters spanned over more than one anatomical or functional region additional peak voxels are reported together with their corresponding anatomical region.

| Cluster | L/R | t-statistic | x | y | z | Voxels |
| --- | --- | --- | --- | --- | --- | --- |
| **A > 0** |  |  |  |  |  |  |
| 1 Heschl's gyrus | L | 12.83 | -48 | -13 | 3 | 7317 |
| Superior temporal gyrus (posterior division) | L | 11.81 | -64 | -16 | 9 |  |

|  |  |  |  |  |  |  |  |
| --- | --- | --- | --- | --- | --- | --- | --- |
|  | Superior temporal gyrus (anterior division) | L | 6.68 | -48 | 12 | -13 |  |
|  | Insula /Operculum | L | 11.2 | -42 | -6 | 18 |  |
|  | Lentiform nucleus (putamen) | L | 4.12 | -31 | -9 | -5 |  |
|  | Lentiform nucleus (lateral globus pallidus) | L | 4.55 | -21 | -13 | -2 |  |
|  | Thalamus (vent.post.lat. nucleus) | L | 3.82 | -17 | -18 | 4 |  |
| 2 | Heschl's gyrus | R | 13.52 | 54 | -9 | 3 | 4046 |
|  | Superior temporal gyrus (posterior division) | R | 12.54 | 64 | -19 | 6 |  |
|  | Superior temporal gyrus (anterior division) | R | 4.32 | 47 | 17 | -9 |  |
|  | Insula | R | 4.16 | 39 | -12 | 20 |  |
| 3 | Splenium of the corpus callosum | L | 4.70 | -19 | -45 | 15 | 194 |
| 4 | Anterior inferior temporal gyrus | L | 4.21 | -36 | -8 | -39 | 165 |
| 5 | Precentral gyrus | R | 4.57 | 57 | -6 | 44 | 32 |
| 6 | Genu of the corpus callosum | L | 3.80 | -17 | 25 | 17 | 28 |
| 7 | Crus cerebri cerebellar peduncles | R | 4.07 | 14 | -15 | -15 | 20 |
| 8 | MTG | L | 3.67 | -62 | -44 | -12 | 20 |
| 9 | IFG | L | 3.38 | -44 | 6 | 23 | 17 |
| 10 | Body of the corpus callosum | R | 3.47 | 2 | 3 | 21 | 15 |

Also, in the left hemisphere we found a cluster in lentiform nucleus with a center in the globus pallidus extending laterally into the putamen and nearby in the ventral posterior lateral nucleus of the thalamus. Smaller clusters were found in the bilateral middle temporal gyrus and the left lingual gyrus as well as the left cerebellar hemisphere and the vermis. We also found clusters in cerebral white matter in the genu and splenium of the corpus callosum at the borders of the anterior and posterior horn of the left ventricle and the body of the corpus callosum at the midline. Finally, we found a small cluster of activation in the right crus cerebri of the cerebellar peduncles. Upon close inspection these white matter clusters did not appear to be the result of “spill over” from nearby grey matter regions. Finally, BOLD activity in the primary visual cortex in this condition was significantly below baseline

### Visual alone (V)

In line with our expectations, the visual alone stimulation resulted in a strong BOLD response in primary visual cortices of bilateral occipital poles (Figure 2, Table 2). The clusters extended laterally to form two prominent foci of activation in the lateral occipital cortices (LOC). From the left LOC region, significant BOLD activity appeared to follow along the ventral visual pathway into the fusiform gyrus and dorsal visual pathway toward the occipito-temporal cortex passing visual motion area MT and MST and extending into the pSTS/G. In the left hemisphere significant BOLD effects of the ventral pathway did not extend anteriorly as far as in the right hemisphere and spared the posterior temporal cortex but showed a small cluster in the pSTS/G. Ventral aspects of the ATLs in both hemispheres showed significant responses but only the left hemisphere also showed clusters in the middle and superior ATL.

**Figure 2.**
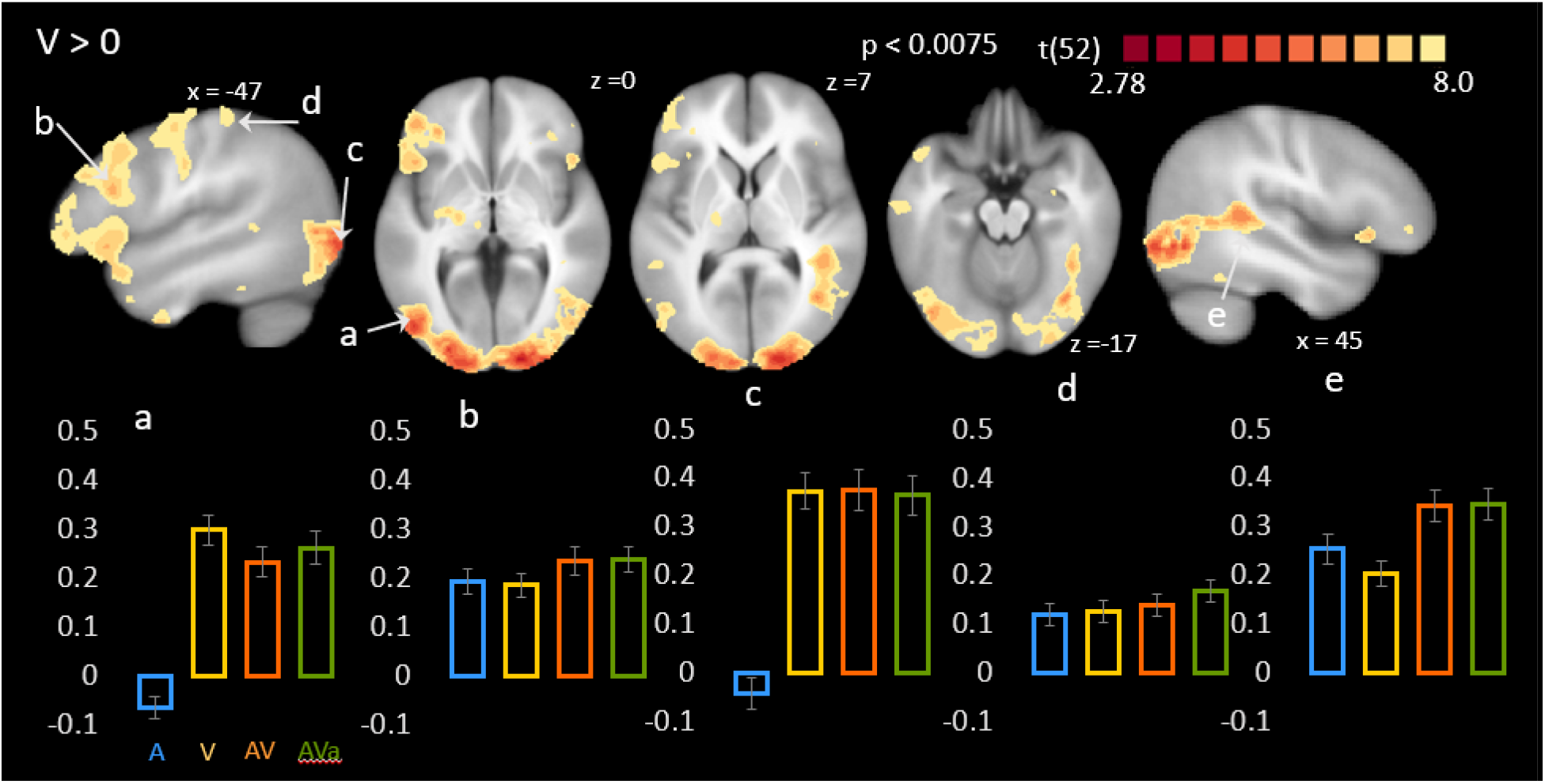
Statistical comparison of the V condition to baseline. Maps show voxels with significant t-scores of the comparison of the V-condition to baseline FDR-corrected (q = 0.05) for multiple comparisons. Bar graphs represent selected % transformed predictor values for A, V, AV and AVa conditions averaged over 4 functional voxels centered around peak voxel locations (see Table 2). a) Left LOC; b) left IFG; c) left occipital pole; d) left precentral gyrus; e) right pSTS.

**Table 2.** Clusters of significant activity resulting from the contrast of the V condition vs. baseline. Significant clusters are numbered and reported with their t-statistic and location in Talairach space in the order of cluster size. In cases where clusters spanned over more than one anatomical or functional region additional peak voxels are reported together with their corresponding anatomical region.

| **V > 0** |  |  |  |  |  |  |  |
| --- | --- | --- | --- | --- | --- | --- | --- |
| 1 | Occipital pole/cuneus | L | 9.12 | -20 | -95 | 2 | 8548 |
|  | Occipital pole/cuneus | R | 9.27 | 17 | -95 | 0 |  |
|  | Lateral occipital complex | L | 8.98 | -43 | -75 | -5 |  |
|  | Lateral occipital complex | R | 6.76 | 45 | -63 | -5 |  |
|  | pSTS | R | 5.47 | 42 | -37 | 8 |  |
| 2 | IFG | L | 5.96 | -58 | 15 | 22 | 3036 |
|  | Precentral gyrus | L | 4.72 | -53 | -5 | 43 |  |
| 3 | Lentiform nucleus (lateral globus pallidus) | L | 4.27 | -21 | -12 | -2 | 265 |
| 4 | MTG | L | 4.36 | -63 | -15 | -13 | 97 |
| 5 | Ventral ATL | R | 4.34 | 37 | -9 | -33 | 89 |
| 6 | Medial frontal gyrus | L | 3.83 | -9 | 33 | 30 | 76 |
| 7 | Amygdala | R | 3.41 | 28 | -1 | -12 | 51 |
| 8 | Pre- and Postcentral gyrus | L | 3.77 | -48 | -29 | 51 | 49 |
| 9 | White matter lateral ventricle | L | 3.45 | -19 | -45 | 16 | 46 |
| 10 | IFG | R | 4.8 | 47 | 21 | -1 | 43 |
| 11 | Medial frontal gyrus | L | 4.24 | -15 | 41 | 45 | 41 |
| 12 | ventral ATL | L | 4.02 | -50 | -3 | -35 | 41 |
| 13 | pSTS/G | L | 3.30 | -51 | -39 | 7 | 38 |
| 14 | Precentral gyrus | R | 4.02 | 57 | -6 | 42 | 35 |
| 15 | Parahippocampus | L | 3.81 | -42 | -29 | -13 | 35 |
| 16 | Culmen | L | 3.49 | -4 | -45 | -9 | 28 |
| 17 | IFG | R | 3.31 | 46 | 39 | 3 | 21 |
| 18 | Thalamus | L | 3.31 | -8 | -18 | 1 | 19 |
| 19 | IFG | R | 3.53 | 34 | 34 | 0 | 16 |
| 20 | Anterior MTG | L | 3.41 | -47 | 13 | -24 | 16 |
| 21 | Precentral gyrus | R | 3.34 | 50 | -9 | 30 | 15 |

There was widespread significant activation in the left frontal lobe involving the ventrolateral prefrontal and lateral frontopolar cortex along the IFG, the nearby frontal operculum and the medial frontal gyrus near the midline. We also found significant clusters in the dlpfc, ventral premotor and ventral and dorsal motor regions. The activations in the right frontal lobe were smaller than in the left hemisphere and included the IFG and lateral frontopolar cortex and primary motor regions. Like in the A>0 contrast, we found a cluster in the left lentiform nucleus and thalamus.

Finally, we found significant activity in the right amygdala and the cerebellar vermis. As in the A > 0 contrast we found activity in the white matter of the splenium of the corpus callosum at the border to the left lateral ventricle.

### MS enhancement: Max. criterion [(AV-A) ⍰ (AV-V)]

The purpose of this conjunction analysis was to identify regions showing MS enhancement where the BOLD response to the AV condition was greater than to the auditory and visual condition respectively (Max criterion). We limited this analysis to regions where AV was greater than baseline (Figure 3, Table 3).

**Figure 3.**
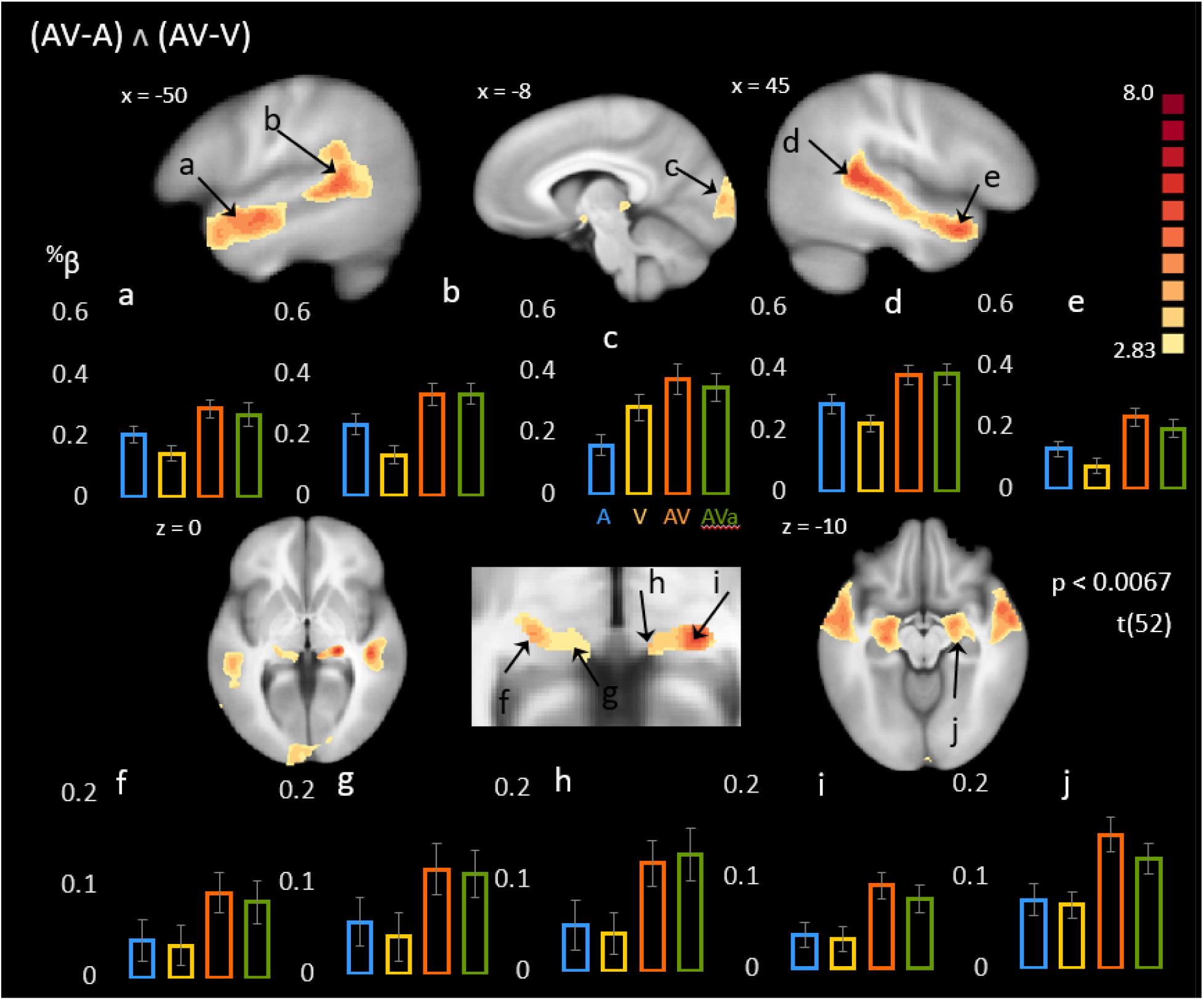
Statistical Conjunction of the (AV-A) and the (AV-V) contrasts (Max. criterion) Maps show voxels with significant t-scores of the conjunction of the (AV-A) and (AV-V) contrasts FDR-corrected (q = 0.05) for multiple comparisons. Bar graphs represent selected % transformed predictor values for A, V, AV and AVa conditions averaged over 4 functional voxels centered around peak voxel locations (see Table 5). a) Left anterior superior temporal gyrus; b) left pSTS; c) left occipital pole; d) right pSTS; e) left anterior STS; f) left LGN; g) left MGN; h) right MGN; i) right LGN; j) right amygdala.

**Table 3.** Clusters of significant activity (Max. criterion) resulting from the conjunction between the AV-A and AV-V contrasts. Significant clusters are numbered and reported with their t-statistic and location in Talairach space in the order of cluster size. In cases where clusters spanned over more than one anatomical or functional region additional peak voxels are reported together with their corresponding anatomical region.

| (AV-A) ^ (AV-V) |  |  |  |  |  |  |  |
| --- | --- | --- | --- | --- | --- | --- | --- |
| 1 | Thalamus (LGN) | R | 6.57 | 21 | -24 | 0 | 3231 |
|  | Thalamus (MGN) | R | 5.936 | 9 | -27 | 1 |  |
|  | Amygdala | R | 4.88 | 21 | -5 | -11 |  |
|  | Thalamus (LGN) | L | 4.64 | -22 | -23 | -2 |  |
|  | Thalamus (MGN) | L | 3.24 | -9 | -26 | -1 |  |
|  | Amygdala | L | 5.34 | -20 | -5 | -13 |  |
| 2 | pSTS | R | 6.74 | 44 | -37 | 8 | 2300 |
|  | Anterior STS | R | 5.81 | 46 | 12 | -18 |  |
| 3 | pSTS | L | 6.31 | -49 | -39 | 6 | 1158 |
|  | Anterior STS | L | 4.81 | -49 | 15 | -18 |  |
| 4 | Cuneus/ occipital pole | L | 4.36 | -5 | -93 | 7 | 566 |
| 5 | Precentral gyrus | R | 3.41 | 53 | -2 | 46 | 28 |
| 6 | IFG | L | 3.36 | -60 | 19 | 20 | 23 |
| 7 | Lingual gyrus | R | 3.33 | 16 | -84 | -1 | 22 |
| 8 | IFG | R | 3.34 | 51 | 22 | 16 | 17 |

We found large MS activations along the STS in both hemispheres spanning from the ATLs into the posterior STS. The posterior sections of these two large clusters extend dorsally to cover the posterior STG and the supramarginal gyrus. While these activations are represented by continuous clusters, they are likely to represent functionally distinct regions.

We therefore increased the statistical threshold in a stepwise fashion to identify local peak activations within the larger STS clusters. We found that both STS clusters contained an anterior, medial and posterior peak in both hemispheres and within the supramarginal gyrus in the left hemisphere. We found significant MS gain in ventral parts of the left temporal lobe and particularly in the bilateral amygdalae. Further, and surprising to us, were bilateral twin clusters in the posterior thalamus encompassing the medial geniculate nuclei, lateral geniculate nuclei and the pulvinar. The conjunction was also significant in several regions of the frontal cortex. These included two smaller clusters in the bilateral IFG and one in the right precentral gyrus (premotor cortex). Finally, the statistical conjunction of audiovisual enhancements over unisensory activations also revealed a significant engagement of the bilateral occipital poles.

### MS enhancement: Superadditivity AV > (A+V)

Here, we examined the distribution of superadditivity in regions where the AV condition was above baseline (Figure 4, Table 4). The FDR-corrected map shows significant superadditivity within the bilateral STS encompassing primary and secondary auditory cortices and more anterior in the STS reaching into the ATL in the right hemisphere. The extracted % transformed beta weights show that in the primary auditory cortices, the AV condition does not significantly exceed the A-condition. Superadditivity is merely due to the V-condition being below baseline. We also found superadditivity within a small cluster of voxels in the left occipital pole and the right supramarginal gyrus in the parietal cortex. Most remarkable, however, was that both MGN clusters survived the statistical threshold, displaying significant effects. We did not find evidence for superadditivity in the posterior STS.

**Figure 4.**
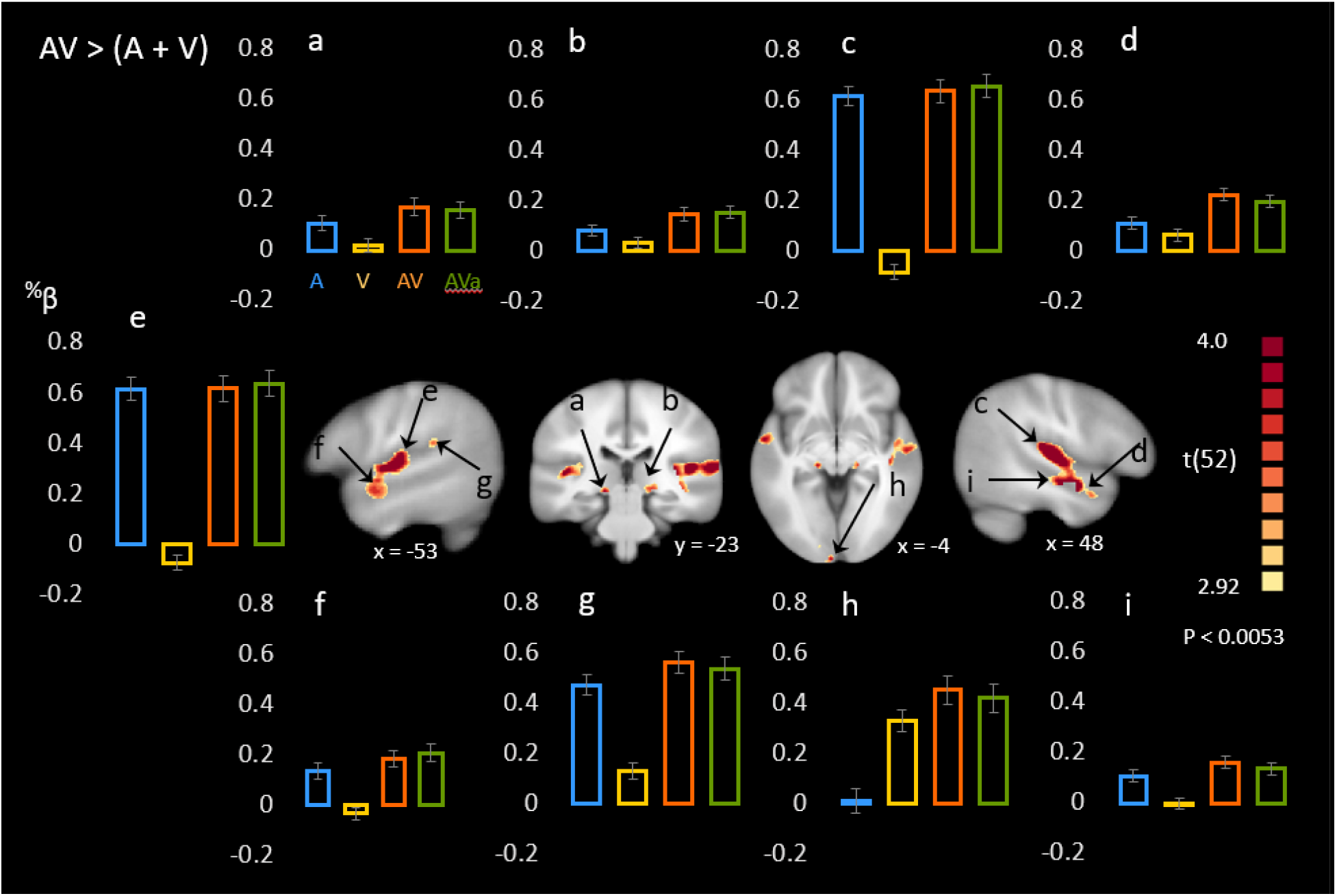
Statistical comparison between AV and (A + V) (Superadditivity) Maps show voxels with significant t-scores of the comparison between the sum of the predictor values of the A and V conditions (A+V) and the AV condition FDR-corrected (q = 0.05) for multiple comparisons fulfilling the superadditive criterion. Bar graphs represent selected % transformed predictor values for A, V, AV and AVa conditions averaged over 4 functional voxels centered around peak voxel locations (see Table 6). a) Left MGN; b) right MGN; c) right pSTS; d) right anterior STS; e) left Heschl’s gyrus; f) left anterior STS g) left pSTS; h) right occipital pole; i) right STS.

**Table 4.** Clusters of significant activity resulting from the subtraction of the sum of the predictor values for the A and V conditions from the AV condition (superadditivity). Significant clusters are numbered and reported with their t-statistic and location in Talairach space in the order of cluster size. In cases where clusters spanned over more than one anatomical or functional region additional peak voxels are reported together with their corresponding anatomical region.

**AV > (A+V)**
|  |  |  |  |  |  |  |  |
| --- | --- | --- | --- | --- | --- | --- | --- |
| 1 | Heschl's gyrus | R | 5.92 | 58 | -11 | 7 | 1244 |
|  | Superior temporal sulcus | R | 5.89 | 52 | -9 | -10 |  |
| 2 | Heschl's gyrus | L | 4.94 | -54 | -15 | 9 | 574 |
|  | Superior temporal sulcus | L | 4.2 | -58 | -3 | -5 |  |
| 3 | Insula (Wernicke) | L | 3.86 | -50 | -34 | 19 | 27 |
| 4 | Lingual gyrus (occipital pole) | L | 3.90 | -5 | -98 | -3 | 26 |
| 5 | STG (temporal pole) | R | 3.67 | 42 | 18 | -19 | 24 |
| 6 | Inf. Occipital gyrus | L | 3.43 | -21 | -89 | -9 | 24 |
| 7 | Medial geniculate nucleus | R | 3.54 | 17 | -23 | -3 | 22 |
| 8 | Medial geniculate nucleus | L | 4.1 | -15 | -23 | -4 | 22 |

### Audiovisual (AV) versus asynchronous audiovisual (AVa)

We found three smaller clusters in which BOLD was higher in the asynchronous condition, one in the right parietal cortex, one in the right prefrontal cortex and one in the left premotor cortex (Figure 5, Table 5). Closer inspection of the betas revealed that for both loci in the right hemisphere both conditions were below baseline. We did not find evidence for relatively increased BOLD activity for the synchronous condition. We therefore concluded that this experimental manipulation did not result in significant differences that are interpretable in regard to our hypotheses.

**Figure 5.**
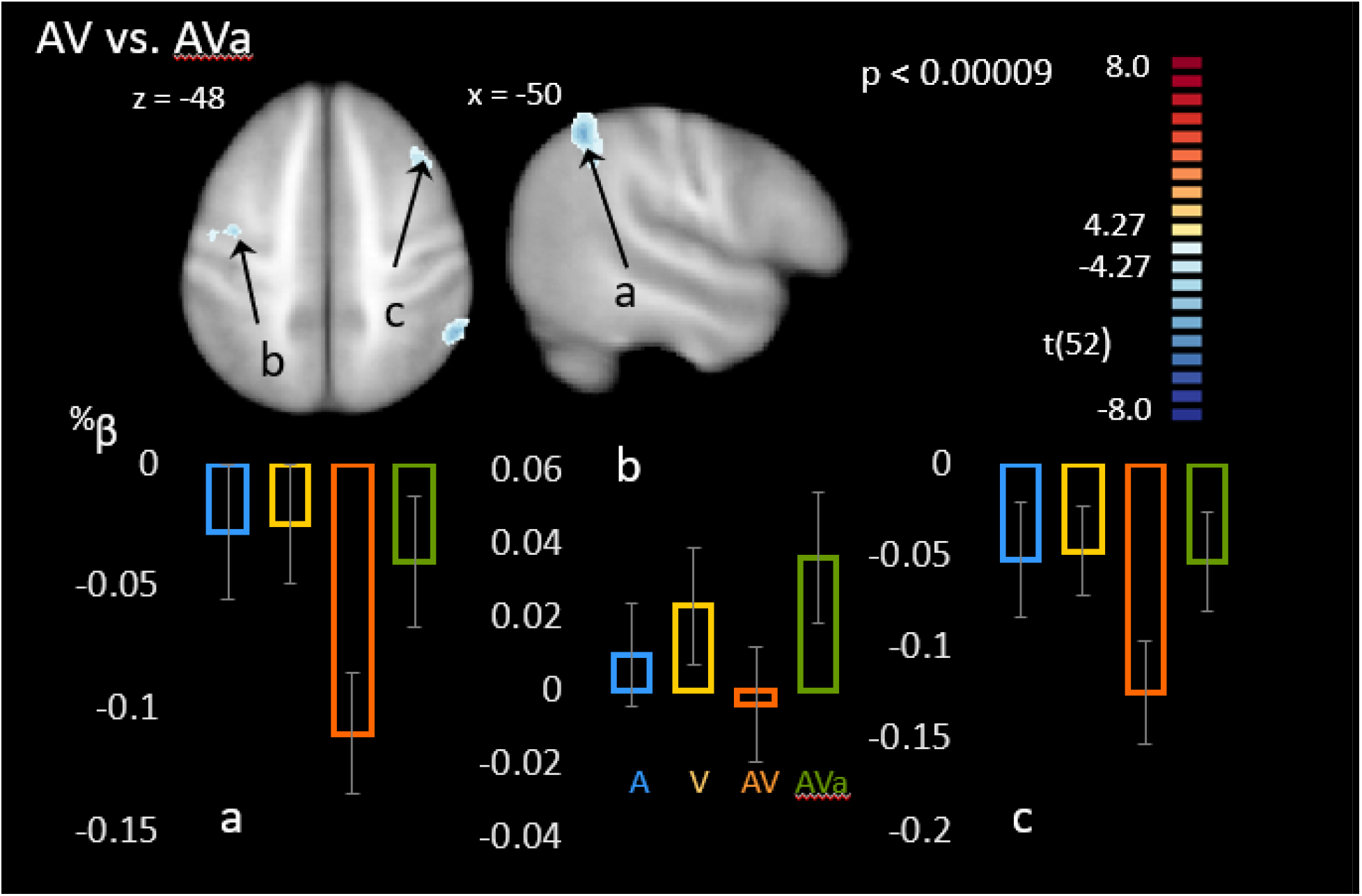
Statistical comparison between AV and Ava conditions. Maps show voxels with significant t-scores of the comparison of the AV and AVa conditions FDR-corrected (q = 0.05) for multiple comparisons. Bar graphs represent selected % transformed predictor values for A, V, AV and AVa conditions averaged over 4 functional voxels centered around peak voxel locations (see Table 7). a) Right parietal lobe; b) left SFG; c) left middle frontal gyrus.

**Table 5.** Clusters of significant differences between the synchronous (AV) and asynchronous (Ava) audiovisual conditions.

| **AV vs. AVa** |  |  |  |  |  |  |  |
| --- | --- | --- | --- | --- | --- | --- | --- |
| 1 | Parietal lobe | R | -6.24 | 52 | -48 | 45 | 203 |
| 2 | Middle frontal gyrus | L | -5.26 | -36 | -8 | 47 | 51 |
| 3 | Superior frontal gyrus | R | -5.22 | 37 | 20 | 48 | 50 |

### A - V

As described in the methods section, we used this contrast as an inclusion criterion for our sample because it was a more sensitive indicator of BOLD responses to the A and V stimuli than comparing unisensory responses to baseline. We also considered that under the likely assumption that in our experimental design the lack of rest conditions between blocks of stimulation could result in a “high” baseline, some unisensory effects would not be observable when comparing to baseline alone.

As expected, the respective A and V conditions resulted in much of the same regional activations they did when compared to baseline (Appendix, Figure 1, Table 1). However, several differences are worth mentioning here. First, and to our initial surprise given the results from the A alone analysis, the A condition resulted in significantly higher BOLD response in much of the visual association cortex with centers of gravity in the bilateral lingual gyrus. This cluster extended dorsally into the parietal cortex around the midline and included the postcentral gyrus. A look at the percent signal change from an ROI at the centers of gravity revealed that all conditions containing a visual stimulus but not the A-condition were significantly below baseline at these locations.

Another remarkable observation was that this contrast revealed activations along the subcortical auditory pathway. The BOLD response in the bilateral medial geniculate nuclei to the A condition was significantly larger than in the V-condition (and larger than any of the conditions containing a visual stimulus). The same pattern was observed in both inferior colliculi although we interpret this with caution due to the small size of these structures and the inability to achieve perfect anatomical matching between subjects.

We also found two clusters (A > V) in the bilateral crus cerebri.

### Conjunction of auditory and visual conditions (A ⍰ V)

We conducted a conjunction analysis to determine which regions of the brain exhibited BOLD responses that were significantly above baseline for both auditory and visual conditions (Appendix, Figure 2, Table 2). We found this to be the case in the right pSTS/G and the ATL, IFG and precentral gyrus in the left hemisphere. We also found the left lentiform nucleus to exhibit a significant response to unisensory A and V stimuli.

### Behavioral task results

In line with previous findings (Ross et al., 2011) (Sumby, 1954) the RM-ANOVA returned main effects (Greenhouse-Geisser corrected for violation of sphericity) of SNR [*F*_(4.78, 239)_ = 16.7; *p* < 0.001; *η*^2^= .25] and condition [*F*_(1, 50)_ = 8.93; *p* = 0.004; *η*^2^= .152] showing that performance decreased as SNR decreased and was significantly better when visualized speech was present (see Figure 6). Audiovisual gain showed the characteristic inverted-u shape relationship to SNR that we have reported in the past (Foxe et al., 2015; Ma et al., 2009; Ross et al., 2015; Ross et al., 2011; Ross, Saint-Amour, Leavitt, Javitt, et al., 2007; Ross, Saint-Amour, Leavitt, Molholm, et al., 2007) with a maximum (*M* = 37.12%; *SD* = 18.7%) at intermediate (−9dB) intelligibility. We found no significant interaction between both factors indicating that SNR affected performance in the A and AV condition in a similar manner [*F*_(4.74, 237)_ = 1.72; *p* = 0.136; *η*^2^= .03]. Neither age [*F*_(1, 50)_ = 3.66; *p* = 0.061; *η*^2^= .068] nor biological sex [*F*_(1, 50)_ = 0.014; *p* = 0.907; *η*^2^< 0.001] were significant. Overall, participants were able to speechread the correct word in *M* = 13.65% (*SD* = 9.61) of cases with no appreciable difference between males (*M* = 12.57%; *SD* = 8.09%) and females (*M* = 14.86%; *SD* = 11.13%) (*F*_(1, 50)_ = 0.516; *p* = 0.476; *η*^2^= 0.01) and no effect of age (*F*_(1, 50)_ = 0.274; *p* = 0.603; *η*^2^= 0.005). We found no relationship between performance in the auditory condition at low SNRs with speechreading performance *r*(51) = 0.013, p = 0.926 and a positive relationship with AV performance at low SNRs *r*(51) = 0.33, *p* = 0.015.

**Figure 6.**
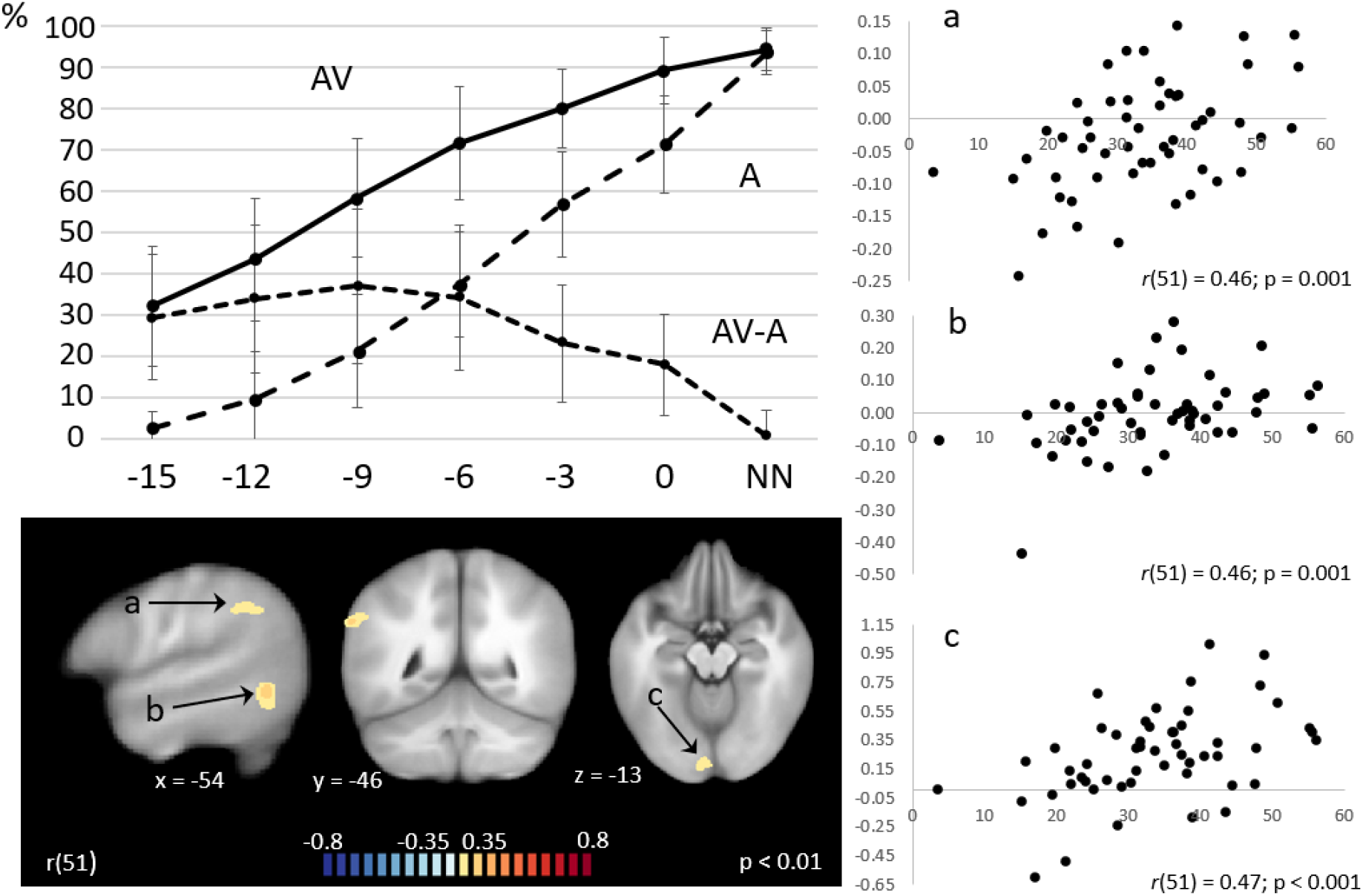
Performance in the behavioral task and associations with brain activity. Line graph represents % correct performance in the A and AV conditions as well as audiovisual gain (AV-A) over seven SNR conditions with error bars representing standard deviations from the mean. Maps represent significant Pearson r correlation coefficients with an applied cluster correction of 74 functional voxels with clusters in the a) cuneus; b) pMTG and c) occipital cortex of the left hemisphere. Scatter plots show % transformed predictor values (AV-A) for each participant averaged over 4 voxels at the centers of the clusters shown in the statistical map in relationship to behavioral audiovisual gain (AV-A) on the x-axis.

### Association between BOLD responses and behavioral performance

#### Audiovisual gain

This analysis was performed to identify brain regions that are involved in the gain conferred by the presence of congruous visual input (Figure 6). For this we conducted voxel-wise correlations between beta weights of the AV-A contrast and the difference between audiovisual (AV) and auditory alone (A) performances in the behavioral experiment. We used the map of the voxel-wise *Pearson r* statistic of the AV-A contrast, thresholded at *p* = 0.01 as the input for the Monte Carlo cluster estimation which, after 5000 iterations returned a cluster threshold of 74 voxels. The resulting map showed left hemispheric clusters of significant positive correlations in the primary visual cortex (*r*(51) = 0.47; *p* < 0.001), the cuneus (*r*(51) = 0.46; *p* = 0.001) and the posterior middle temporal gyrus (*r*(51) = 0.46; p = 0.001).

#### Audiovisual

Cluster threshold estimation was carried out on the r-map reflecting significant correlations between the AV BOLD response and AV performance in the SIN task thresholded at *p* = 0.01. Only a cluster in the inferior parietal lobe survived a threshold of 67 voxels (*r*(51) = 0.36; *p* = 0.008).

#### Visual alone (Speechreading)

The correlation map between BOLD response to the V-condition and performance in the speechreading condition was thresholded at *p* = 0.01 and submitted to the cluster threshold estimation resulting in a minimum cluster size of 69 voxels. No cluster in the map survived this threshold.

#### Auditory alone

We computed whole brain voxel-wise correlations between the BOLD predictor of the A-condition and the average performance (% correct) in the auditory condition of the speech in noise behavioral task. The correlation map was thresholded at *p* = 0.01 and submitted to the Monte Carlo cluster estimation procedure which returned a cluster threshold of 62 voxels. No plausible significant correlations between the BOLD response and behavioral performance were found (a cluster in the cerebellum was driven by two outliers).

#### Questionnaire performance

Forty eight out of 53 participants completed the 10-item questionnaire which can be found in the appendix. A majority of seventy five percent of the participants answered six or more questions correctly and the average number of correct answers was 7.6 (*SD* = 2.2). Due to the interspersed V-alone blocks, it was fully expected that task performance would remain below a perfect score. Note that this experiment was designed with an eye towards future investigations of MSI processes across development and was therefore constructed to be suitable for use in children (hence the choice of a narrative that would appeal to all age groups). The presentation of a continuous narrative precluded the use of a simultaneous behavioral task, so our intention here was to ensure task compliance via our instruction that the subject would be “tested” after the scan. We included the five adults for whom we did not have the questionnaire data because 1), eye-tracking measures in these individuals made it clear that they fixated the screen appropriately with eyes open throughout the experiment, and 2), we inspected the statistical maps for each subject to ensure the presence of typical auditory and visual sensory activation patterns indicating compliance with experimenter instructions.

#### Correlation with BOLD measures

After cluster threshold estimation (*p* = 0.001) voxel-wise correlation map between questionnaire performance and the A > 0 contrast was further constrained to clusters larger than 27 functional voxels. We found a significant cluster in the left STS *r*(46) = 0.51; *p* < 0.001 (X: −44, Y: −12, z: −7; 544 voxels) (see Figure 7). We also found a significant correlation between questionnaire performance and the AV condition at the same location. Since both conditions are highly correlated its report is omitted here.

**Figure 7.**
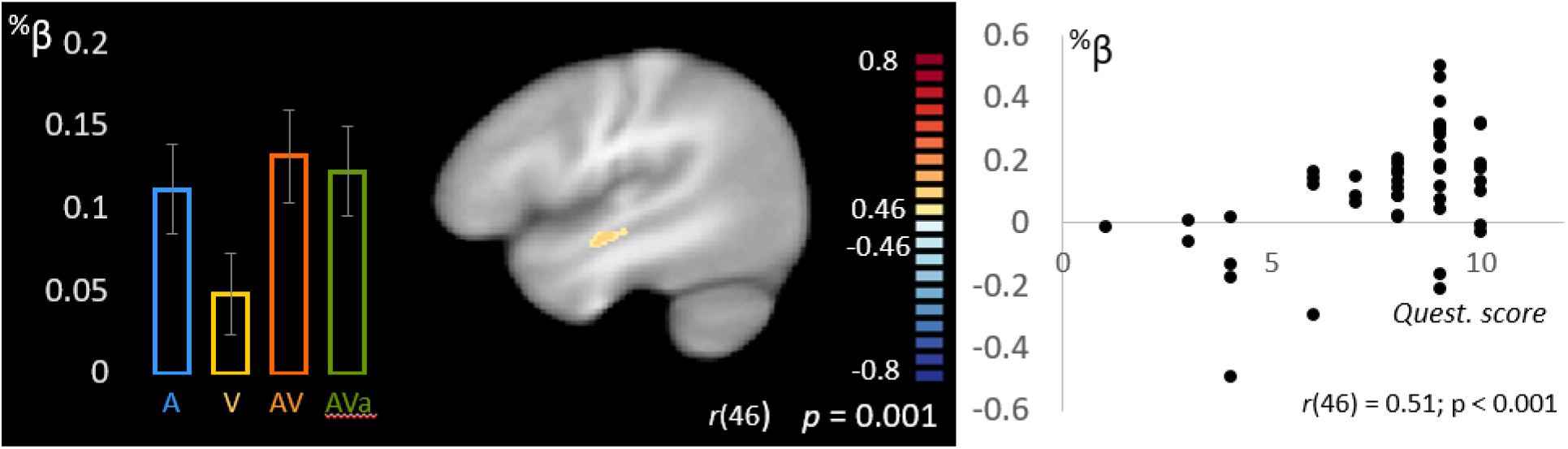
Association between questionnaire performance and brain activity (A condition) The map shows voxels with significant Pearson correlation coefficients in 48 subjects of the association between questionnaire performance and predictor values in the A condition at a cluster correction threshold of 27 voxels. Bar graphs represent % transformed predictor values for A, V, AV and AVa conditions averaged over 4 functional voxels centered around the peak voxel location of the cluster in the left STS. The cluster plot shows the relationship between questionnaire performance (x-axis) and %-transformed predictor values in the A condition averaged over 4 voxels centered around the peak location of the cluster in the left STS.

## DISCUSSION

The goal of this fMRI study was to investigate brain regions showing audiovisual enhancement during perception of narrative speech. We expected this enhancement to be evident in regions previously identified as part of the MS speech network (Erickson et al., 2014) (Calvert & Thesen, 2004) but also in regions downstream from known MSI sites reflecting the effects of successful integration rather than MSI alone.

### Unisensory conditions

Before we assessed MS enhancement, we explored how the unisensory narrative speech stimulus engaged the speech network. At the word level, core perisylvian language areas are active (Binder, 1997; Binder et al., 2000; Hertrich et al., 2020). Sentences and narratives differ in several regards from more simple speech stimuli, such as words or phonemes that have been used in most previous studies of audiovisual speech processing, and are known to involve core perisylvian language regions. The addition of syntactic and semantic information at the sentence level involves additional perisylvian cortex “spreading” along the posterior STS/G into the ATLs (Ardila et al., 2016; Binder, 2017; Price, 2012). Narratives contain additional, more complex semantic information in the form of thematic content tying information delivered over multiple sentences into a common overarching context (Hertrich et al., 2020; Xu et al., 2005; Xu et al., 2017). Further, processing at discourse level requires the listener to extract meaning increasingly tying lexical information to world knowledge creating mental representations of the narrative (Xu et al., 2005) and their potential social implications. This places additional demands on attention, working memory and higher cognitive functions such as theory of mind and may evoke emotions and visual imagery. Therefore, the increasing complexity of the language stimulus involves a multitude of extralinguistic cognitive operations involving extrasylvian regions that are reflected in the BOLD signal (de Heer et al., 2017; Huth et al., 2016; Lerner et al., 2011). This increase in complexity has been shown to engage left frontal regions (Lerner et al., 2011) supported by evidence from studies in frontotemporal dementia (Ash et al., 2006; Peelle & Grossman, 2008).

Our results support these findings and extend them in several respects. We found bilateral activation of perisylvian regions along the superior temporal plane and bilateral engagement of the articulatory motor and supplementary motor cortex but activity in the IFG and dlPFC was left lateralized.

We found it interesting that the V condition (V > 0) appeared to engage an extended network of left frontal regions including the IFG. One possible explanation is that participants subvocalize during speechreading or are engaged in cognitive processes that evoke semantic activity. On the other hand, the contrast between the unisensory conditions (A vs. V) did not result in higher activations to the A condition in these same regions as one would expect, since these frontal regions are involved in operations resulting from narrative speech processing (Binder et al., 2009; Hertrich et al., 2020; Xu et al., 2005).

We made several further novel observations when contrasting the A and V conditions directly (A vs. V). First, it was apparent that the bilateral MGNs are engaged during listening to natural narrative speech. This is expected, given that their function as thalamic relays along the auditory pathway is well known. However, effects in these subcortical structures are rare in fMRI experiments due to their small size and accompanying issues that will be discussed further below.

We also observed that activation in much of the visual association cortex was larger to the auditory alone stimulus than to the visual articulation, especially in the bilateral lingual gyrus. All three conditions containing a visual stimulus are far below baseline whereas the auditory stimulus was at baseline. This pattern is reversed in the nearby primary visual cortex at the occipital pole. We speculate that activity in visual association cortex to the auditory stimulus is due to visual imagery evoked by the story narrative that is presumably absent in the V condition. We further speculate that this activity is suppressed in AV conditions when a visual stimulus is present because of a shift of attention to the visual stimulus (i.e. the speaker). This is an incidental finding not related to the original purpose of the study but is nevertheless interesting and relevant to report because it may shed light on a mechanism related to evoked visual imagery during auditory stimulation and its suppression.

### Audiovisual enhancement

We used the following conjunction approach [(AV-A) ⍰ (AV-V)] to identify regions of MS enhancement. This was largely true for regions along the left and right STS and included posterior sections of the STS commonly associated with MSI (Beauchamp et al., 2004).

The STS is well known to be part of the semantic system (Binder et al., 2009; de Heer et al., 2017; Hickok et al., 2018) and these findings provoke the question why these regions are enhanced by a MS stimulus. One explanation could be that despite our efforts to make the stimulus intelligible under unisensory auditory stimulation, the additional information from visual articulation resulted in an increase in intelligibility which in turn affected the content processed by the semantic system. If this was the case, however, this increase in intelligibility would likely be evident in modality specific auditory regions in the superior temporal plane. We did not find evidence for a significantly higher BOLD effect in the AV condition than in the A condition due to possible ceiling effects in these regions (also see discussion on superadditivity). Responses in the STS to the unisensory conditions were overall lower, leaving “room” for MS enhancement.

Another possible explanation is that the MS stimulus is inherently more salient than unisensory stimuli alone. Moreover, more than just articulatory features used for linguistic analysis, it conveys important non-linguistic contextual information through tone, timing and volume of the voice, facial expressions, posture and head movement (Munhall & Buchan, 2004; Munhall & Johnson, 2012). In a natural conversation this additional information may be used by the speaker to aid the delivery of information, clarify intent and project emotional state. In the case of the reading of a story by a trained actress, as was the case in our experiment, this MS contextual information is more complex because the speaker does not deliver her own state or intent but that of the characters and their roles in the story. Given the complexity of a natural narrative, it is apparent how this non-linguistic contextual information renders the MS stimulus particularly salient. The notion of a widespread, non-specific effect of saliency of the MS stimulus is supported by our finding of MS enhancement in the bilateral amygdalae. Neither visual speech articulation and emotional facial expression nor listening to the auditory narrative with its emotional content was sufficient to engage the amygdalae compared to baseline.

MS enhancement was also observed in the primary visual cortex around the occipital poles. Less prevalent but significant activity was also found in higher-order areas considered part of the ventral visual pathway in inferior temporal regions including the TFC and the anterior ITG of the left hemisphere. These regions correspond well with regions previously identified as involved in visual speech perception (Bernstein & Liebenthal, 2014). Thus, MS speech integration is not simply associated with visual influences on auditory processing, but rather, there is a clear bi-directionality to these influences, with substantial modulation of visual processing seen as a result of auditory inputs. This corresponds well with findings from a human intracranial study by our group where auditory inputs significantly impacted visual cortical processing of simultaneously presented visual inputs across a substantial extent of visual cortex (Mercier et al., 2013).

### Subcortical audiovisual enhancement

Perhaps the most striking finding of this study is that of focal enhancements in the posterior thalamus involving the medial and lateral geniculate nuclei and the pulvinar. That subcortical structures such as the superior colliculus (Wallace et al., 1998; Xu et al., 2014; Yu et al., 2013), inferior colliculus (Gruters & Groh, 2012) and some of the thalamic nuclei, especially the medial pulvinar (Cappe et al., 2009; Dietrich et al., 2013; Froesel et al., 2021), are involved in MS processing is now well-known. However, we typically think of these structures as involved in integration of relatively simple stimuli and it might be considered somewhat surprising to find that thalamic nuclei are playing a prominent role in audio-visual speech processing (although see (Hebb & Ojemann, 2013)).

A spate of neuroimaging studies has indeed pointed to such subcortical MS processing under a variety of conditions. For example, Noesselt and colleagues (Noesselt et al., 2010) showed that functional connectivity between both the medial and lateral geniculate nuclei with their respective sensory cortices, as well as with the STS, was modulated under MS conditions and that the strength of these couplings across participants was associated with performance on a visual stimulus detection task for difficult-to-detect low-contrast visual inputs. Using a MS target detection task where synchronized auditory “pips” have been found to substantially improve target detection in cluttered moving visual scenes (Van der Burg et al., 2008), van der Burg and colleagues asked if variance in this MS ability could be associated with both structural and functional connectivity between thalamic nuclei and sensory-cortical representations. Using diffusion tensor imaging and probabilistic tractographic techniques, they asked whether connectivity between task-specific auditory and visual cortex (A1 and V4) and in turn, between these regions and their respective thalamic nuclei, would predict inter-individual differences in MS target detection. They found that the strength of structural connectivity between the cochlear nucleus, the medial geniculate body and primary auditory cortex was related to this integrative ability.

Perhaps more directly relevant to the current work is evidence from studies using speech stimuli showing MS responses in the brainstem. Fairhall and Macaluso (Fairhall & Macaluso, 2009) showed that attention to congruent AV speech stimuli resulted in increased activation in the superior colliculus compared to attention to incongruent stimuli. In an electrophysiological study, Musacchia et al. (Musacchia et al., 2006) recorded the auditory brainstem response (ABR) while participants listened to synthesized phonemic stimuli (e.g. /da/). There were three different conditions, one with no visual input where only the phonemes were heard, one where phonemes were accompanied by either congruent visual articulations or incongruent visual articulations, and one where only the visual tokens were presented during silence. They found modulation of both latency and amplitude of the auditory brainstem response (ABR) under audio-visual conditions, effects that began as early as 11 ms following acoustic input, and these MS effects were also found to differ as a function of congruence between the visual and acoustic phonemic inputs. The work suggests, as do our results here, that ongoing visual articulatory inputs can shape the auditory system’s response to anticipated acoustic inputs, and that this top-down modulatory effect can be instantiated extremely early in the subcortical processing hierarchy – indeed, even before auditory information reaches the relevant thalamic nuclei. It is of interest to note that recordings directly in rat auditory thalamus have shown that visual inputs can substantially modulate the early phase of auditory thalamic responsivity, significantly impacting behavior in these animals (Komura et al., 2005).

### Superadditivity

The comparison (AV > A + V) in fMRI is largely adopted from animal electrophysiology studies that have shown neurons exhibiting stronger responses to MS as opposed to unisensory stimulation (Stein & Stanford, 2008; Xu et al., 2014). The rationale for adopting this method for BOLD fMRI that reflects activity from large populations of neurons was that BOLD activation is a time invariant-linear system where activation to two stimuli presented together is equivalent to the sum of the two stimuli presented individually (see James (James, 2012) for a review). If a region contains MS cells, the evoked activity to a MS stimulus is predicted to exceed the sum of the unisensory responses.

In practice this theoretical model to identify MS regions has proven to be too conservative, with many fMRI studies failing to show superadditivity in regions well known to be involved in MSI (Beauchamp, 2005; James, 2012). Since the explicit goal of the present study was to explore a network of regions showing MS enhancement beyond the sites typically reported to be involved in MS integration *per se*, we adopted the less conservative max criterion (Altieri et al., 2011; Beauchamp, 2005). We conducted an additional analysis of MS enhancement using the additive criterion with the goal to explore whether this criterion would isolate a reduced set of classic MS integration sites, expecting these regions to overlap with clusters identified with the max criterion.

The results of this analysis highlighted some of the problems associated with this criterion. First, we failed to replicate superadditivity in the posterior STS. In the temporal lobes, superadditivity is apparent in Heschl’s gyrus and the superior temporal plane in both hemispheres covering large parts of the auditory cortex but only in the more anterior STS. In the auditory cortex, the effect is mainly due to the fact that V is below baseline. The difference between AV and A is not significant. If one assumes that brain activity in the auditory cortex as measured as BOLD effects reflects the quality of the perceptual effect then it would be hard to argue that the AV condition would convey any perceptual advantage over the A condition. We attribute the lack of difference between the A and AV conditions to the high intelligibility of the auditory stimulus with the result of a ceiling effect.

Therefore, in conditions of high intelligibility and considering the somewhat arbitrary nature of the baseline, superadditive effects can be misleading. Nevertheless, we found genuine superadditive effects in the bilateral MGNs, the anterior portions of the STS and a small cluster in the left occipital pole.

### MS temporal congruency

One way to overcome the inherent difficulty of identifying MS regions by comparing MS to unisensory BOLD responses (James, 2012; Stevenson et al., 2009) is to adopt an experimental approach that allows for comparison of two MS conditions to one another that engage MS regions differentially. The approach adopted here that was successfully used in the past (Miller & D’Esposito, 2005; Stevenson et al., 2010; van Atteveldt et al., 2007; van Wassenhove et al., 2007) was to offset the auditory and the visual tracks sufficiently to prevent an integration of sound and visual articulatory movements. Based on the extant literature, a 400 ms delay of the visual signal appeared appropriate for this purpose. Participants are able to detect an asynchrony of an audiovisual speech signal at a 132ms delay of the visual signal (Dixon, 1980) see also (van Wassenhove et al., 2007). The strength of the McGurk effect is reliably different at a 60ms delay of the visual signal (Munhall & Buchan, 2004; Munhall et al., 1996). In an fMRI study by Stephenson et al. (Stevenson et al., 2010), a 400 ms offset (visual lead) was effective in generating BOLD differences between synchronous and asynchronous audiovisual stimulus material.

Based on previous findings (Marchant et al., 2012; Noesselt et al., 2007; Stevenson et al., 2010) (Okada et al., 2013)we expected the temporal congruency of the auditory and visual speech inputs to impact the degree to which the MS network was engaged, particularly in the posterior superior temporal cortex.

Much to our surprise, our experimental manipulation was not effective in evoking the expected effects. One reason could be that an offset of 400ms was not sufficient to prevent integration from taking place or that over the course of the experiment, participants managed to adapt to the asynchrony, perhaps as a result of entrainment unfolding over the course of the experiment (Crosse et al., 2015; Luo et al., 2010). Another possible explanation is that in the asynchronous condition groups of MS neurons are engaged despite the lack of synchronicity and thus drive a BOLD response that is comparable to the synchronous AV condition. The activity of this group of neurons may not reflect a response to congruous auditory and visual information and would not result in MS enhancement under more degraded listening conditions. Since our stimuli were sufficiently intelligible, this activity may have had no detrimental effect on the perception of the auditory stimulus and therefore did not result in a difference in the BOLD signal between MS conditions.

### Correlation with behavioral multisensory speech-in-noise task

The motivation for this analysis was to locate regions associated with performance in an audiovisual speech perception task. However, we would like to advise the reader to consider the results of this particular analysis as well as their interpretation as preliminary. According to a publication by Eklund et al., (Eklund et al., 2016) we were not able to meet sufficiently conservative criteria for the protection against false positives because the initial correlation maps before cluster threshold estimation did not exceed *p* < 0.001 for whole brain analysis (see methods section). We expected these regions to be part of the well-established perisylvian speech processing network. However, it was in fact the primary visual cortex, cuneus and the posterior middle temporal gyrus (pMTG) of the left hemisphere that showed significant association with MS gain in the behavioral task. The involvement of the primary (V1) and secondary (cuneus) visual cortex suggests that the ability to benefit from visual articulation is associated with activity in the visual cortices and may reflect processes underlying the analysis of visual articulatory movement and/or attention to the visual stimulus (Vanni et al., 2001). There is strong evidence that the pMTG plays a key role in semantic cognition. Davey et al. (Davey et al., 2016) suggested that this structure integrates information related to more automatic aspects of semantic cognition (presumably associated with passive listening) often associated with the default mode network and information processing associated with effortful task-related semantic retrieval. This model fits well with the notion that during passive listening to a complex narrative in our experiment, semantic aspects of the DMN and task-related semantic retrieval are both engaged depending on the degree of effort required to follow the thematic content of the story. While we ensured that the auditory stimulus in our experiment was sufficiently intelligible, it is likely that intelligibility varied over the course of the experiment and/or between subjects due to the difficulty to fully control this variable in a scanner environment causing a stimulus-dependent, flexible change of engagement of passive default mode and effortful task-dependent semantic retrieval. The ability to engage the pMTG might in turn be related to the ability to retrieve semantic information in the low-intelligibility context of our speech-in-noise experiment and therefore serve as a possible explanation for the correlation between pMTG BOLD signal and MS gain in the behavioral experiment. Finally, despite its intuitive appeal, we did not find support for the notion that individuals with difficulty perceiving degraded speech exhibit greater speechreading or greater audiovisual benefit under difficult listening conditions. In fact, we found a positive association between A and AV conditions at low SNRs, the opposite of what is predicted under this hypothesis.

We did find an association between questionnaire performance and brain activation to the A and AV conditions which emerged in regions of the STS where we also found significant audiovisual enhancement, a region generally considered to support semantic processing (de Heer et al., 2017; Hertrich et al., 2020; Hickok & Poeppel, 2007) which is likely to be modulated by the complexity and salience of the language stimulus (Ross & Olson, 2010; Xu et al., 2005). The most parsimonious explanation for this effect would simply be that subjects with lesser engagement of the semantic network were less likely to answer questions about the semantic content of the narrative correctly. However, this could have been caused in several ways such as lack of attention to the stimulus, quality of the stimulus delivery (although differences in brain activation at sensory/perceptual levels were not apparent in our data) or differences in cognitive/semantic processing itself. Further, due to the interspersed V-alone conditions, some subjects might have missed critical aspects for the understanding of the full narrative which could have caused both questionnaire performance and lower brain activity.

### Conclusion

The current study, by using a naturalistic narrative stimulus set and imaging a substantially larger cohort than used in most previous studies, revealed a considerably more extensive network of MS enhancement. This network included “classic” sites of MS integration as well as parts of the semantic language network. We also found enhancement in extralinguistic regions not usually associated with MS integration, namely the primary visual cortex and the bilateral amygdalae. Analysis also revealed involvement of thalamic brain regions along the visual and auditory pathways more commonly associated with early sensory processing

We posit that under natural listening conditions, MS enhancement not only involves sites of MS integration but many regions of the wider semantic network and includes regions associated with extralinguistic perceptual and cognitive processing.

## List of abbreviations

ATL: anterior temporal lobe
AV: audiovisual condition
AVa: asynchronous audiovisual condition
IFG: inferior frontal gyrus
Dlpfc: dorsolateral prefrontal cortex
MS: multisensory
MSI: Multisensory integration
MTG: middle temporal gyrus
pMTG: posterior medial temporal gyrus
STG: superior temporal gyrus
pSTS/G: posterior superior temporal sulcus/gyrus

## Appendix

**Figure 1.**
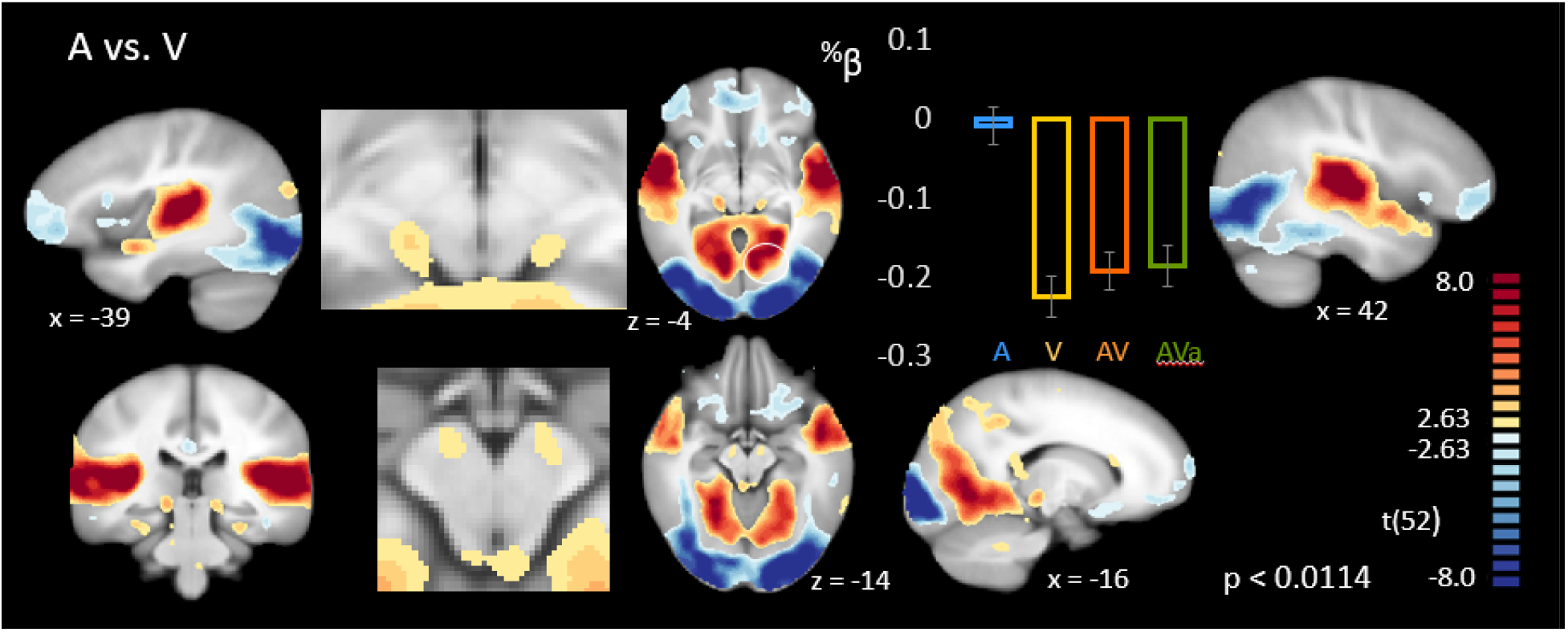
Statistical comparison of A and V conditions. Maps show voxels with significant t-scores of the comparison of the A and V conditions FDR-corrected (q = 0.05) for multiple comparisons. The bar graphs represents selected % transformed predictor values for A, V, AV and AVa conditions averaged over 4 functional voxels centered around the peak voxel location (A > V) in the cuneus/ lingual gyrus (circled) (see Table 3).

**Table 1.** Clusters of significant activity resulting from the contrast between A and V conditions. Significant clusters are numbered and reported with their t-statistic and location in Talairach space in the order of cluster size. In cases where clusters spanned over more than one anatomical or functional region additional peak voxels are reported together with their corresponding anatomical region.

| A – V |  |  |  |  |  |  |  |
| --- | --- | --- | --- | --- | --- | --- | --- |
| A > V |  |  |  |  |  |  |  |
| 1 | Cuneus/lingual gyrus | R | 10.60 | 7 | -64 | 9 | 17364 |
|  | Cuneus/lingual gyrus | L | 9.31 | -3 | -74 | 17 |  |
| 2 | Heschl's gyrus | L | 17.56 | -47 | -18 | 7 | 7425 |
| 3 | STG | R | 16.62 | 54 | -8 | 2 | 5861 |
| 4 | Middle frontal gyrus | R | 3.15 | 21 | -2 | 57 | 85 |
| 5 | Cerebellar tonsil | L | 3.40 | -30 | -42 | -32 | 74 |
| 6 | Cerebellar tonsil | L | 3.24 | -15 | -44 | -33 | 60 |
| 7 | Pyramis of cerebellar vermis | L | 3.36 | -4 | -75 | -29 | 56 |
| 8 | Cerebellar peduncle | R | 4.04 | 11 | -14 | -15 | 55 |
| 9 | Genu of the corpus callosum | L | 3.27 | -16 | 23 | 17 | 38 |
| 10 | Corpus callosum body | L | 3.27 | -1 | -1 | 24 | 38 |
| 11 | Medulla | R | 3.11 | 1 | -29 | -39 | 35 |
| 12 | MTG | L | 3.23 | -64 | -43 | -11 | 23 |
| 13 | Cerebellar peduncle | L | 3.48 | -9 | -12 | -13 | 21 |
| 14 | Splenium of the corpus callosum | R | 3.14 | 2 | -32 | 13 | 19 |
| *V > A* |  |  |  |  |  |  |  |
| 1 | Middle occipital gyrus | R | -13.10 | 26 | -91 | 0 | 12109 |
| 2 | Superior frontal gyrus (BA 10) | L | -5.24 | -23 | 59 | 4 | 4224 |
| 3 | Insula | L | -4.60 | -31 | 18 | 1 | 747 |
| 4 | Insula | R | -4.77 | 36 | 20 | 3 | 556 |
| 5 | Medial frontal gyrus | L | -4.13 | -1 | 30 | 34 | 385 |
| 6 | Inferior parietal lobe | L | -3.33 | -49 | -54 | 41 | 227 |
| 7 | Superior frontal gyrus (BA 6) | L | -4.30 | -2 | 4 | 57 | 121 |
| 8 | Superior frontal gyrus (BA 9) | L | -3.71 | -20 | 45 | 34 | 83 |
| 9 | Caudate nucleus | L | -3.94 | -7 | 1 | 12 | 81 |
| 10 | Cingulate gyrus | L | -3.34 | -2 | -25 | 28 | 49 |
| 11 | Superior frontal gyrus | L | -3.08 | -20 | 38 | 44 | 37 |
| 12 | Supramarginal gyrus | R | -2.98 | 57 | -51 | 27 | 34 |
| 13 | MTG | L | -3.82 | -55 | -30 | -12 | 30 |
| 14 | Cerebellum (inf. semilunar lobule) | R | -3.08 | 43 | -66 | -35 | 28 |
| 15 | Superior frontal gyrus (BA 8) | L | -3.04 | -5 | 16 | 48 | 23 |
| 16 | Cingulate gyrus | R | -3.59 | 2 | 13 | 26 | 23 |
| 17 | Middle frontal gyrus (BA 8) | L | -3.05 | -45 | 25 | 38 | 21 |

**Figure 2.**
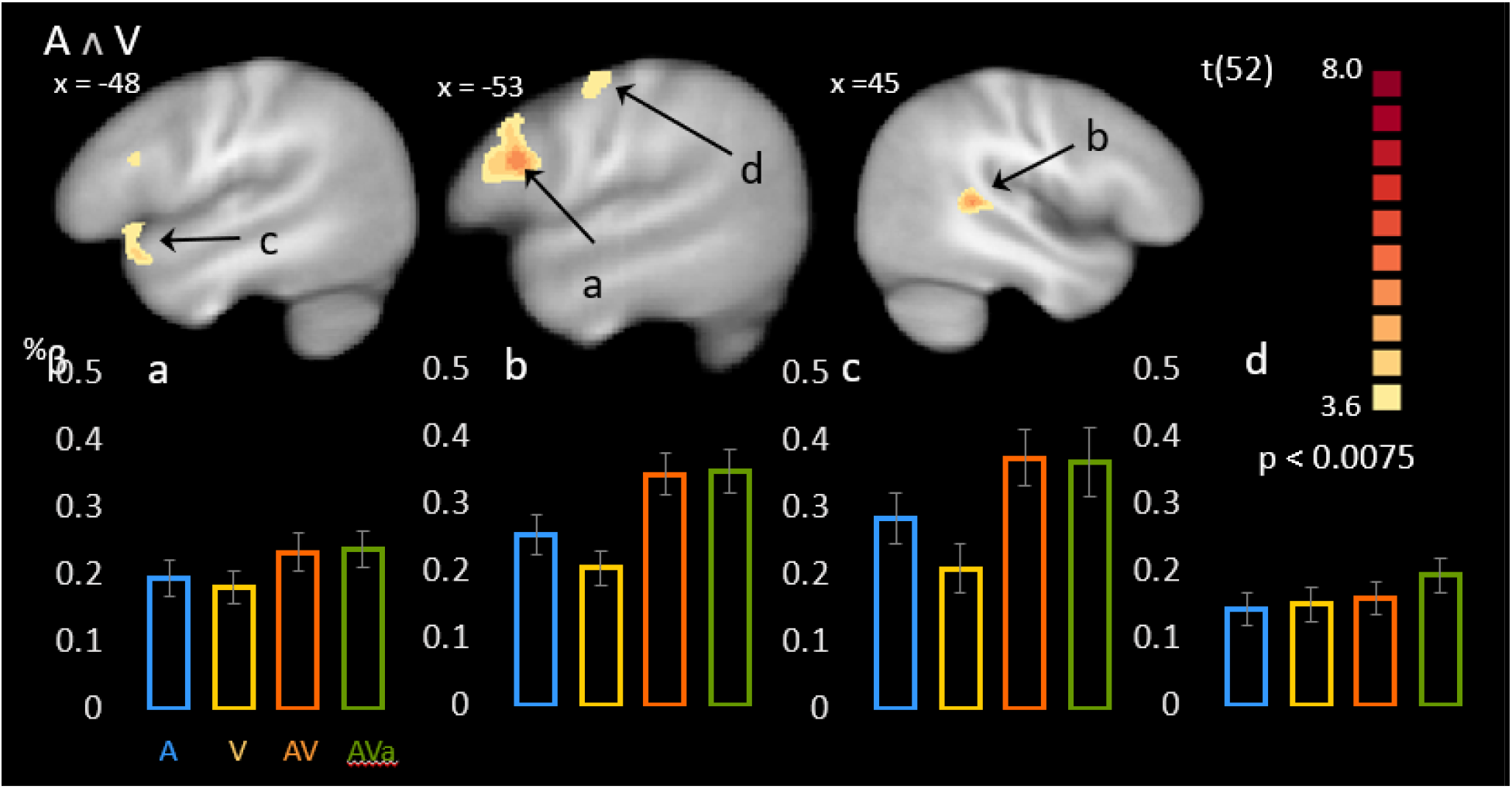
Statistical Conjunction between A and V conditions. Maps show voxels with significant t-scores of the conjunction analysis of the A-condition to baseline FDR-corrected (q = 0.05) for multiple comparisons. Bar graphs represent selected % transformed predictor values for A, V, AV and AVa conditions averaged over 4 functional voxels centered around peak voxel locations (see Table 1). A) Left IFG; B) right pSTS; C) left anterior STS; D) left precentral gyrus.

**Table 2.** Clusters of significant activity resulting from the conjunction between the A vs. baseline and V vs. baseline contrasts. Significant clusters are numbered and reported with their t-statistic and location in Talairach space in the order of cluster size.

|  |  |  |  |  |  |  |  |
| --- | --- | --- | --- | --- | --- | --- | --- |
| 1 | IFG | L | 5.81 | -57 | 14 | 21 | 230 |
| 2 | Posterior superior temporal gyrus | R | 5.23 | 43 | -38 | 8 | 116 |
| 3 | Anterior superior temporal gyrus | L | 4.18 | -48 | 18 | -12 | 64 |
| 4 | Precentral gyrus | L | 4.15 | -54 | -7 | 43 | 42 |

## Notes

### Competing Interest Statement

The authors have declared no competing interest.

